# Advancing apple genetics research: *Malus coronaria* and *Malus ioensis* genomes and a gene family-based pangenome of native North American apples

**DOI:** 10.1101/2024.05.13.593769

**Authors:** Anže Švara, Honghe Sun, Zhangjun Fei, Awais Khan

## Abstract

Wild *Malus* species flourished in North America long before Europeans introduced domesticated apples. *Malus coronaria* and *M. ioensis* are native to the mid-western and eastern USA, while *M. angustifolia* and *M. fusca* grow in the southeast and west, respectively. They offer disease resistance, climate and soil adaptability, and horticultural traits for apple breeding. However, their utilization remains limited due to insufficient genomic resources and specific genetics. We report high-quality phased chromosome-scale assemblies of *M. coronaria* and *M. ioensis*, generated using long-read and conformation capture sequencing. Phylogenetic and synteny analysis indicated high relatedness between these two genomes and previously-published genome of *M. angustifolia*, and lower relatedness with *M. fusca*. Gene family-based pangenome of North American *Malus* identified 60,211 orthogroups containing 340,087 genes. Genes involved in basic cellular and metabolic processes, growth, and development were core to the existence of these species, whereas genes involved in secondary metabolism, stress response, and interactions with other organisms were accessory and are likely associated with adaptation to specific environments. Structural variation hotspots were mostly overlapping with high gene density. This study offers novel native North American *Malus* genome resources that can be used to identify genes for apple breeding and understand their evolution and adaptation.

## Introduction

Wild *Malus* species could provide novel genes and alleles to broadening the genetic base of apple cultivars [1]. Four small-fruited wild *Malus* species including the Southern (*M. angustifolia*), Sweet (*M. coronaria*), Oregon/Pacific (*M. fusca*), and Prairie (*M. ioensis*) crabapples originate from four distant geographic regions across the North America and are reputed for their phenotypic and genetic distinctiveness from domesticated apples [2]. For example, late blooming *M. angustifolia* is adapted to relatively high temperatures [3], *M. coronaria* is capable of producing seed through asexual (apomixis) and sexual reproduction [4], *M. fusca* is a source of fire blight resistance [5], and *M. ioensis* accessions are known for their ornamental qualities [6]. Ancestors of *M. fusca* are believed to arrive to North America by the Bering Strait [7], and have subsequently diversified from other three North American species [2]. The evolutionary diversification of North American *Malus* species requires further genomic clarification [2].

Genomes of apple species are characterized by complexity due to large size, high heterozygosity and sometimes by polyploidy as a result of the open pollination and evolutionary hybridizations. Out of four North American species, *M. fusca* and *M. angustifolia* genomes have been assembled recently [3,5]. The haplome assemblies have size of ∼635 Mb and ∼717 Mb, respectively, and 0.8% and 1.1% estimated heterozygosity. In contrast to minimal structural divergence between *M. domestica* cultivars and the Eurasian *Malus* progenitors, the North American genomes showed larger diversification and structural variation. *M. angustifolia* genome emerged as the most distinct species, compared to other *Malus* species [3].

Unique characteristics and heterozygous regions of individual genome assembly offer opportunities to compare genomic regions that are associated with phenotypic variation in a pangenome [8]. Reference *Malus* pangenomes were generated to capture genetic diversity in terms of SNPs and indels by resequencing numerous accessions, revealing the domestication and wild introgression history of domesticated apple and their primary progenitor species *M. sieversii* and *M. sylvestris* [9]. Thousands of genes originate from the progenitors, of which hundreds are fixed in cultivated apple through hybridization [9]. Allele-specific expression of genes is associated with traits such as fruit quality [9], and different gene copy-number variation across the genomes [10]. Studies have shown that sequencing multiple reference genomes can better represent genetic diversity and relatedness of the whole population, detect structural variants (SVs), and pinpoint candidate genes associated with a trait [8,9].

Pangenome of a group of accessions is defined as a sum of core, accessory, and specific genomes of a group of more or less related individual organisms [11]. Core genomes typically anchor genes critical for common traits of the group that are present in all the tested accessions. They can vary in size in a given species or genus and can consist of ∼30-90% of all the identified genes [12]. Core genes in *Malus* pangenome of three species contained 81.3–87.3% of all genes, which is more than in perennial shrubs and annual plants such as cranberry (*Vaccinium macrocarpon* Aiton; 53%), blueberry (*Vaccinium* spp. L.; 51%), rice (*Oryza sativa*; 80%), corn (*Zea mays*; 40%) [9,12–14]. Core genomes typically harbor genes enriched for ‘housekeeping’ functions such as basal metabolism and circadian cycle [10,12]. In contrast, accessory and specific genomes are mostly associated with adaptive functions vital for survival of a more specific groups of organisms in a divergent environment [10,12]. For example, stress response, pathogen defense, specialized metabolism were enriched in dispensable genomes of apples, purple false brome (*Brachypodium distachyon*), and European cabbage (*Brassica oleracea*) [10,12,15,16]. The genes found in accessory and specific genomes are of particular interest for breeders looking for novel traits among a wide variety of accessions [10,12].

Here, we *de novo* assembled and annotated phased chromosome-scale genomes of two native North American *Malus* species, *M. coronaria* and *M. ioensis*. In conjunction with two previously assembled reference genomes of *M. fusca* and *M. angustifolia* we built a phylogeny tree across *Malus* species including Eurasian *Malus* species, and identified syntenic regions across the analyzed genomes. Furthermore, we constructed a gene family-based pangenome containing genic regions of North American *Malus* species. Functional enrichment for core-, accessory, and specific genomes has been explored to pinpoint evolutionary important gene groups and their functions. Structural variation of the coding and noncoding regions has been evaluated for future studies of species-specific traits.

## Materials and methods

### PacBio HiFi and Illumina Hi-C sequencing

Dormant cuttings of one-year-old wood of *M. coronaria* accession PI590014 and *M. ioensis* accession PI590015, both originating from the central USA (Fig. 1A-B), were collected from the research orchard at USDA ARS - Plant Germplasm Resources Unit (PGRU) at Cornell AgriTech in Geneva, NY, USA. The cuttings were placed in cups containing water until young leaf buds emerged in the dark. Once the newly formed shoots with young leaves emerged, they were sampled and immediately frozen at -80°C. The samples were shipped on dry ice to the DNA Sequencing & Genotyping Center at the University of Delaware, USA for high-molecular-weight genomic DNA extraction and Pacific BioSciences (PacBio) Single Molecule Real Time (SMRT) sequencing as previously described by Khan et al. (2022). The extracted DNA was fragmented into 15-kb fragments and HiFi library was constructed by the use of SMRTbell Express Template Prep Kit 2.0 for PacBio sequencing. DNA/Polymerase Binding Kit 2.0 (Pacific Biosciences) was utilized according to the manufacturer’s protocol and the library was filtered for fragments >10 Kb using Sage Blue Pippin (Sage Sciences) to remove smaller fragments and adapter dimers. PacBio Sequel IIe was used in the CCS/HiFi mode with a single SMRT cell with 2 hours of pre-extension and 30-hour movie times to sequence the library. PacBio HiFi reads were filtered for adapters using pacbio.RemoveAdapters2 function in bbmap v39.01 [18]. The read length distribution and quality of the obtained reads were assessed using Pauvre v0.1923 [19] and assembly-stats v1.0.1 [20].

**Figure 1:**
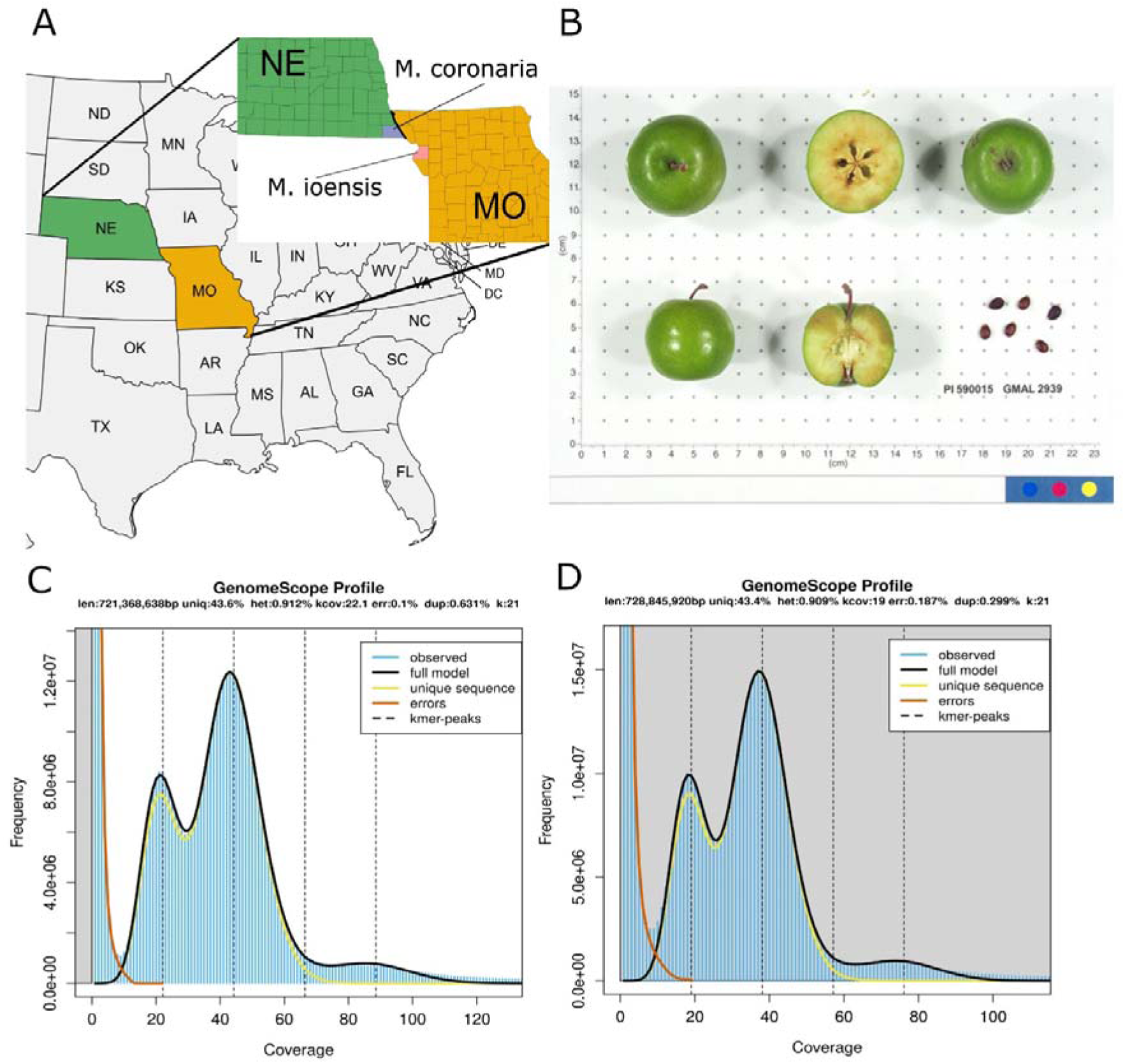
Genome assemblies of for *Malus coronaria* accession PI590014 and *Malus ioensis* accession PI590015. **A)** Collection sites of PI590014 and PI590015 in Nebraska and Missouri, respectively. **B)** *M. ioensis* (PI590015) produces middle-sized green fruits of 3-4 cm in diameter (source: USDA ARS - GRIN Global). **C-D)** k-mer plots of PI590014 (C) and PI590015 (D).

A total amount of 1 g of flash-frozen young leaf material was harvested from the leaves of each accession to extract DNA and perform Illumina Hi-C chromatin conformation capture sequencing at DNA Sequencing & Genotyping Center of the University of Delaware, USA. The library was prepared using the Dovetail Genomics Omni-C kit to be sequenced on an Illumina NovaSeq 6000 with PE150 reads and fastqc v0.12.1 was used to validate the overall quality of the library [21].

### Phased haplome assembly and scaffolding

21-mers were generated using raw HiFi data using Jellyfish v2.3.0 (RRID:SCR_005491) [22] to estimate the genome size and heterozygosity by GenomeScope 2 [23], similarly to our previous work [17,24]. The reads were phased, purged and assembled into contigs by hifiasm v0.16.1 that integrated Hi-C reads as an additional input to facilitate phasing of the contigs without triobining function (RRID:SCR_021069) [25]. Plastid contigs were identified by MitoHiFi v3.0.0 [26] and filtered from the contiguous assembly. The assemblies were evaluated and trimmed for adaptor and foreign organism DNA using NCBI Foreign Contamination Screening (FCS) pipeline fcs 0.3.0, i.e., by using FCS-adaptor and FCS-GX [27].

Scaffolding of contigs into chromosomes was performed by Juicer pipeline v2.0 (RRID:SCR_017226) [28] with parameters ‘-C 20000000 -l regular -s none’ with all the Hi-C reads. 3D-DNA v08012019 was used to generate genomic contact maps by running the run-asm-pipeline.sh function with parameters ‘--mode haploid --editor-coarse-resolution 100000 --editor-coarse-region 400000 --editor-saturation-centile 10 -- editor-repeat-coverage 50’ [29]. Contact maps were visually inspected, manually edited and oriented in Juicebox Assembly Tools (JBAT) v2.17.00 (RRID:SCR_021172) [30] to generate 17 chromosomes of a *Malus* haplome. run-asm-pipeline-post-review.sh with the parameter ‘-g 500’ was used to generate fasta files of the haplomes. The assemblies were aligned with the GDDH13 assembly [31] using MUMmer v4.0 (RRID:SCR_018171) [32] and Assemblytics [33] was used to generate synteny dotplot and to determine the number and orientation of specific chromosomes by inspecting alignment coordinates. Genome quality and completeness were assessed using BUSCO v5.2.2 (RRID:SCR_015008) [34] with the “eudicots_odb10” database. Haplome quality values and k-mer completeness were assessed using Merqury v1.3 [35].

### Repeat annotation and gene prediction

The haplome assemblies of *M. coronaria* and *M. ioensis*, as well as the previously assembled *M. fusca* were annotated following the pipeline described previously [24]. *M. angustifolia* assembly was not included in this analysis, as the published version is hard-masked and is hence unsuitable for annotations. Briefly, the assemblies were masked using repeat sequence libraries previously constructed from the genomes of ‘Gala’, *M. sieversii*, and *M. sylvestris* [9]. Redundant repeat sequences in these libraries were removed using the ‘cleanup_nested.pl’ script in the EDTA package (v 2.1.0) with default parameters [36]. The resulting non-redundant repeat library was used to mask each haplome using RepeatMasker (v4.0.8; http://www.repeatmasker.org/). Protein-coding genes were predicted from the repeat-masked assemblies with MAKER pipeline [37], by integrating evidence from *ab initio* gene prediction, transcript and protein evidence. AUGUSTUS [38] and SNAP [39] were used for *ab initio* gene predictions. Genome-guided transcript assembly of previously reported transcriptomic data and CDS sequences from different apple genomes [9,31,40] were used as transcript evidence, and protein sequences of published apple, peach, strawberry, and Arabidopsis genomes, and the UniProt database (Swiss-Prot plant division) were used as protein homology evidence. To functionally annotate the predicted genes, their protein sequences were searched against the SwissProt and TrEMBL databases (https://www.uniprot.org/) using BLASTP with an e-value cutoff of 1E-5, and the InterPro database (https://www.ebi.ac.uk/interpro/) using InterProScan [41].

### Phylogeny and synteny analyses

BUSCO v5.2.2 (RRID:SCR_015008) [34] with the “eudicots_odb10” database was ran on two concatenated haplomes of newly and previously assembled North American *Malus* species, the assemblies of *M. sieversii*, *M. sylvestris*, and *M. domestica* GDDH13 and ‘Gala’, and *Pyrus communis* L. [42] and *Prunus armeniaca* [43] to build a phylogenetic tree of relatedness between different assemblies [3,5,9,17]. BUSCO_phylogenomics v2023-12-17 [44] that concatenates and aligns single-copy BUSCO genes from each assembly was used to construct the alignment matrix that can be used for phylogeny tree construction. IQ-tree2 v2.2.2.6 [45] was used to build a phylogenetic tree by incorporating 1000 bootstrap replicates to establish node support values and the best-fitting model that was identified by the ModelFinder module [46]. The maximum likelihood tree was plotted in the Interactive Tree Of Life (iTOL) online tool [47].

The newly assembled haplomes and the first haplomes of *M. fusca* and *M. angustifolia* were subjected to a pairwise synteny comparison according to their relatedness obtained by the above phylogeny analysis to identify major structural differences between assemblies. MCscan (Python version) v0.8.12 pipeline [48] was utilized to identify syntenic regions among the assemblies with parameters ‘-m jcvi.compara.catalog ortholog --cscore=0.99 and -m jcvi.compara.synteny screen --minspan=10’. Synteny plots were generated with the parameter ‘-m jcvi.graphics.karyotype’.

### Construction of gene family based pangenome

Gene clusters or orthogroups comprising pan, core, accessory, and specific genes were constructed using OrthoFinder v2.5.4 [49]. Pangenome was defined as the set of orthogroups identified in any of all eight haplomes of the four North American *Malus* species, whereas the core genome comprised the groups shared among these haplomes. Soft-core gene clusters were defined as those present in any combination of seven haplomes, and accessory genome comprised orthogroups found in three to six haplomes, or in two haplomes, when these two haplomes originate from different accessions. Finally, species-specific orthogroups were identified in individual haplome assemblies only or two haplomes originating from the same accession.

The obtained sizes of pan- and core genomes as determined by the number of orthogroups in each individual haplome were utilized to model pan- and core genome sizes. The model was fitted in R using the nonlinear least squares based on a modified version of the Levenberg–Marquardt nlsLM function from package minpack.lm (v1.2-4) [50], as described previously [15]. Points used in regression corresponded to all the possible combinations of genomes, and the pangenome size was modelled using the power law regression y = Ax^B^ + C and the core genome size was modelled using exponential regression y = Ae^Bx^ + C.

Enrichment analysis of core, accessory, and specific genomes was performed by comparing the gene ontology (GO) terms assigned to genes found in the corresponding orthogroups compared to those in the pangenome. GO enrichment analysis was performed using the Singular Enrichment Analysis (SEA) function of the agriGO v2.0 online platform [51] with default settings.

### Haplome alignments and identification of structural variants

Whole-haplome alignments were performed in a reference-based manner using minimap2 v2.27 [52], where each of the newly-assembled haplomes of *M. coronaria*, *M. ioensis*, as well as the haplomes of *M. fusca* were aligned to GDDH13 reference assembly to generate variation data that can be compared across different assemblies [31]. The generated minimap2 output files were fed to SyRI v1.6.3 [53] to identify structural variants (SVs) larger than 50 bp, encompassing insertions, deletions, inversions, and translocations. The distribution of the variation breakpoints was calculated for each 400 kb window along each chromosome. Top 5% of all windows containing the highest abundance of SVs was defined as SV hotspots. The distribution of SVs across the apple genome was visualized using Circos v0.69-9 [54]. Genes overlapping the hotspots were used to perform gene ontology enrichment analysis using Singular Enrichment Analysis (SEA) function of the agriGO v2.0 online platform [51].

## Results

### Haplotype-resolved chromosome-scale assemblies of M. coronaria and M. ioensis

In total, 2,566,711 and 2,122,071 PacBio HiFi reads with an average length of 12,728 and 13,619 bp were generated for *M. coronaria* and *M. ioensis*, respectively, resulting in a total of 32.7 and 28.9 Gb of sequences. These corresponded to ∼45× and ∼40× coverage of the *M. coronaria* and *M. ioensis* genomes that had estimated sizes of 721.4 Mb and 728.8 Mb, and heterozygosity levels of 0.91% and 0.83%, respectively, based on k-mer analysis (Fig. 1C-D). Next, 295 and 265 million paired-end Hi-C reads summing up to 47.3 and 42.6 Gb were generated to facilitate scaffolding of contigs into chromosomes (Fig. S1). *De novo* assembly integrating long and short reads yielded two phased haplomes that were highly contiguous and similar in size (Table 1, Fig. S2). Haplomes 1 (HAP1) and 2 (HAP2) of *M. coronaria* were 666.6 and 675.5 Mb long, and 644.2 and 661.4 Mb in *M. ioensis*. They were scaffolded from 620-923 *M. coronaria* contigs with N50 of 2.3-2.4 Mb, whereas *M. ioensis* haplomes contained 859-1056 contigs with N50 of 2.3 Mb (Table 1). The final haplome assemblies of both genomes contained 17 chromosomes. *M. coronaria* haplomes showed a *k*-mer completeness rate of 97.7% and a consensus quality value (QV) of 67.0 based on the analysis using Merqury [35] (Table 1), whereas *M. ioensis* showed 95.8% k-mer completeness and a QV of 66.2. The BUSCO completeness of *M. coronaria* haplomes was 97.5-98.0% and that of *M. ioensis* was 97.2-98.0% (Table 1, Fig. 2A).

**Figure 2:**
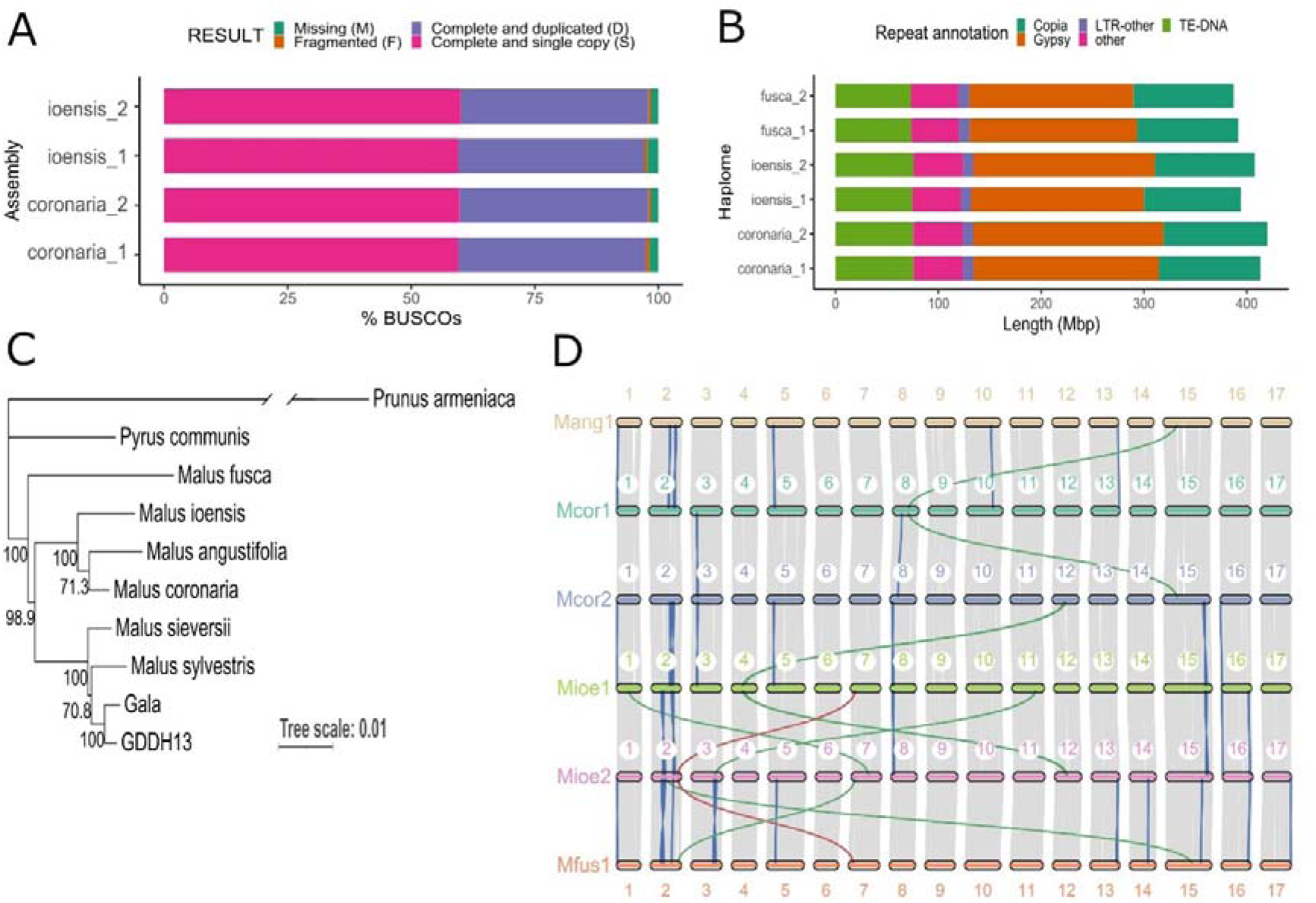
Comparison of *Malus coronaria* and *M. ioensis* haplome assemblies. A) BUSCO completeness of the assembled *M. coronaria* and *M. ioensis* haplomes. B) Repeat components of the North-American *Malus* species including *M. coronaria*, *M. ioensis* and *M. fusca*. C) Phylogenetic analysis of the North-American *Malus* species, *M. domestica* ‘Gala’ and GDDH13, and their primary progenitor species *M. sylvestris* and *M. sieversii* [9,31] using *Pyrus communis* L. [42] and *Prunus armeniaca* [43] as outgroups. Branch of the outgroup *Prunus ameniaca* was broken to fit the plot. D) Macro-synteny plot comparing both haplomes of *M. coronaria* and *M. ioensis* and haplome 1 of *M. angustifolia* and *M. fusca*. Blue, green, and red colors connecting chromosomes indicate inversions, translocations, and inverted translocations, respectively.

**Table 1:** Assembly statistics of *Malus coronaria* (PI590014) and *M. ioensis* (PI590015) haplomes.

| Species | *coronaria* |  |  | *ioensis* |  |  |
| --- | --- | --- | --- | --- | --- | --- |
| Assembly | Haplome 1 | Haplome 2 | Sum | Haplome 1 | Haplome 2 | Sum |
| Total chromosome length (bp) | 666,594,626 | 675,532,332 | 1,342,126,958 | 644,188,171 | 661,368,478 | 1,305,556,649 |
| Contig N50 | 2,321,080 | 2,438,609 |  | 2,314,429 | 2,260,716 |  |
| Gaps | 609 | 589 | 1,198 | 577 | 589 | 1,166 |
| Complete BUSCOs (%) | 97.5 | 98.0 |  | 97.2 | 98.0 |  |
| Quality value | 67.1 | 66.8 | 67.0 | 66.6 | 65.9 | 66.2 |
| k-mer completeness (%) | 83.9 | 84.5 | 97.7 | 81.4 | 82.7 | 95.8 |

### Annotation statistics

Annotation of repeat sequences, and gene prediction of newly assembled *M. coronaria* and *M. ioensis*, and previously assembled *M. fusca* [3,5] genomes enabled direct comparison among the three North-American *Malus* species. In contrast, *M. angustifolia* was hard-masked and was only used for analyses of protein-coding regions. The *M. coronaria*, *M. ioensis*, and *M. fusca* genomes contained between ∼387.2 and ∼420.0 Mb repeats that represented 61.3-62.2% of the assembled genome sizes (Table S1). LTR elements were the most abundant repeats consisting of 42.4-43.9% of the assemblies, followed by DNA transposons that represented 11.2-11.6% of the assemblies (Fig. 2B). Gypsy repeats represented the largest part of the LTRs, occupying over 25% of *M. coronaria*, *M. ioensis*, and *M. fusca* genomes, while copia repeats were present across ∼15% of the three genomes. Other repeats including LINE, SINE, satellite DNA, simple repeats, and other unknown repeat sequences represented less than 10% of the genome assemblies.

The number of annotated genes ranged between 43,773 in haplome 1 of *M*. *coronaria* and 46,068 in haplome 1 of *M*. *fusca*, with an average of 44,755 genes per haploid genome (Table 2). Per gene, 5.3 exons were identified, and the average length of coding sequence (CDS) ranged between 1137.1 bp in haplome 1 of *M. angustifolia* and 1148.0 in *M. ioensis* haplome 2. BUSCO analysis indicated that the annotated genes showed high completeness, ∼98% for all diploid assemblies (Table S3).

**Table 2:** Gene annotation statistics of *Malus coronaria* (PI590014) and *M. ioensis* (PI590015) haplomes, and the annotation of previously-assembled *M. angustifolia* and *M. fusca* haplomes [3,5]

|  | coronaria |  | ioensis |  | angustifolia |  | fusca |  |
| --- | --- | --- | --- | --- | --- | --- | --- | --- |
|  | haplome1 | haplome2 | haplome1 | haplome2 | haplome1 | haplome2 | haplome1 | haplome2 |
| Number of genes | 43,773 | 44,100 | 43,846 | 44,083 | 45,351 | 44,985 | 46,068 | 45,895 |
| Average CDS length | 1143.7 | 1143.4 | 1143.7 | 1148.0 | 1137.1 | 1143.1 | 1137.9 | 1139.6 |
| Average number of exons per gene | 5.3 | 5.3 | 5.3 | 5.3 | 5.3 | 5.3 | 5.3 | 5.3 |
| Average exon length | 214.4 | 212.6 | 213.9 | 213.7 | 213.4 | 213.1 | 215.2 | 215.9 |

### Phylogeny and synteny

Analysis of relatedness between different assemblies based on BUSCO single genes revealed phylogenetic relationships among Eurasian and North American *Malus* species (Fig. 2C). North American species showed distinct grouping from Eurasian species. *M. coronaria* and *M. ioensis* were part of the same clade that encompassed *M. angustifolia* within the central-eastern North American group of *Malus* species. *M. fusca* formed a distinct branch than other North American species. Finally, minimal divergence in genome relatedness was observed between Gala and GDDH13 and their primary Eurasian progenitor species *M. sieversii* as they formed a single clade.

Analysis of the macrosynteny in genic regions at the chromosomal level unveiled extensive synteny and conserved orientation of genic segments of all 17 chromosomes across newly assembled haplomes as well as with *M. angustifolia* and *M. fusca* haplome 1 (Fig. 2D). Only a few macro-structure variations were discovered. Haplomes 1 and 2 of *M. coronaria* contained two inverted regions on chromosome 3 and 8 and a translocation from chromosomes 8 of haplome 1 to chromosome 15 of haplome 2. Haplomes 1 and 2 of *M. ioensis* contained six inversions over four different chromosomes, and four translocations of which one was inverted. Similarly, ten or less inversions and three or less translocations were discovered between *M. coronaria* and *M. ioensis* haplomes, and in their comparison to *M. angustifolia* and *M. fusca* haplome 1.

### Gene family based pangenome composition, modeling, and enrichment

Gene family based pangenome analysis and modeling identified specific orthogroups belonging to pan, core, soft-core, accessory, accession-specific, and haplome-specific genome of North American *Malus* species (Fig. 3). In total, 60,211 orthogroups containing 340,087 genes were identified in the haplomes of *M. coronaria*, *M. ioensis*, *M. angustifolia*, and *M. fusca* (Fig. 3A). From these, the largest group of genes belonged to the core genome, which contained 19,312 orthogroups and 166,477 genes, whereas soft-core genome that encompass all combination of seven haplomes contained 5,786 orthogroups and 45,391 genes. Accessory genome, encompassing 2-6 haplomes, contained 26,198 orthogroups and 108,627 genes. The lowest number of orthogroups and genes was found in accession- and haplome-specific genomes, which contained 6,936 and 1,979 orthogroups, respectively, and 14,706 and 4,886 genes. The number of genes identified in each of the group represent 49.0%, 13.3%, 31.9%, 4.3, and 1.4% of the pangenome genes, respectively. Pan- and core genome model based on the eight assemblies showed that the pangenome is increasing in size with the addition of new genomes, whereas the core genome size is decreasing (Fig. 3B). The respective increase and decrease in genome size were the most pronounced at the level of one to two genomes, where the change in size contained 7,405 and 7,915 orthogroups. The addition of new genomes resulted in the lowest change in the number of orthogroups from seven to eight genomes, where the difference was only 1,458 and 253 orthogroups, respectively.

**Figure 3:**
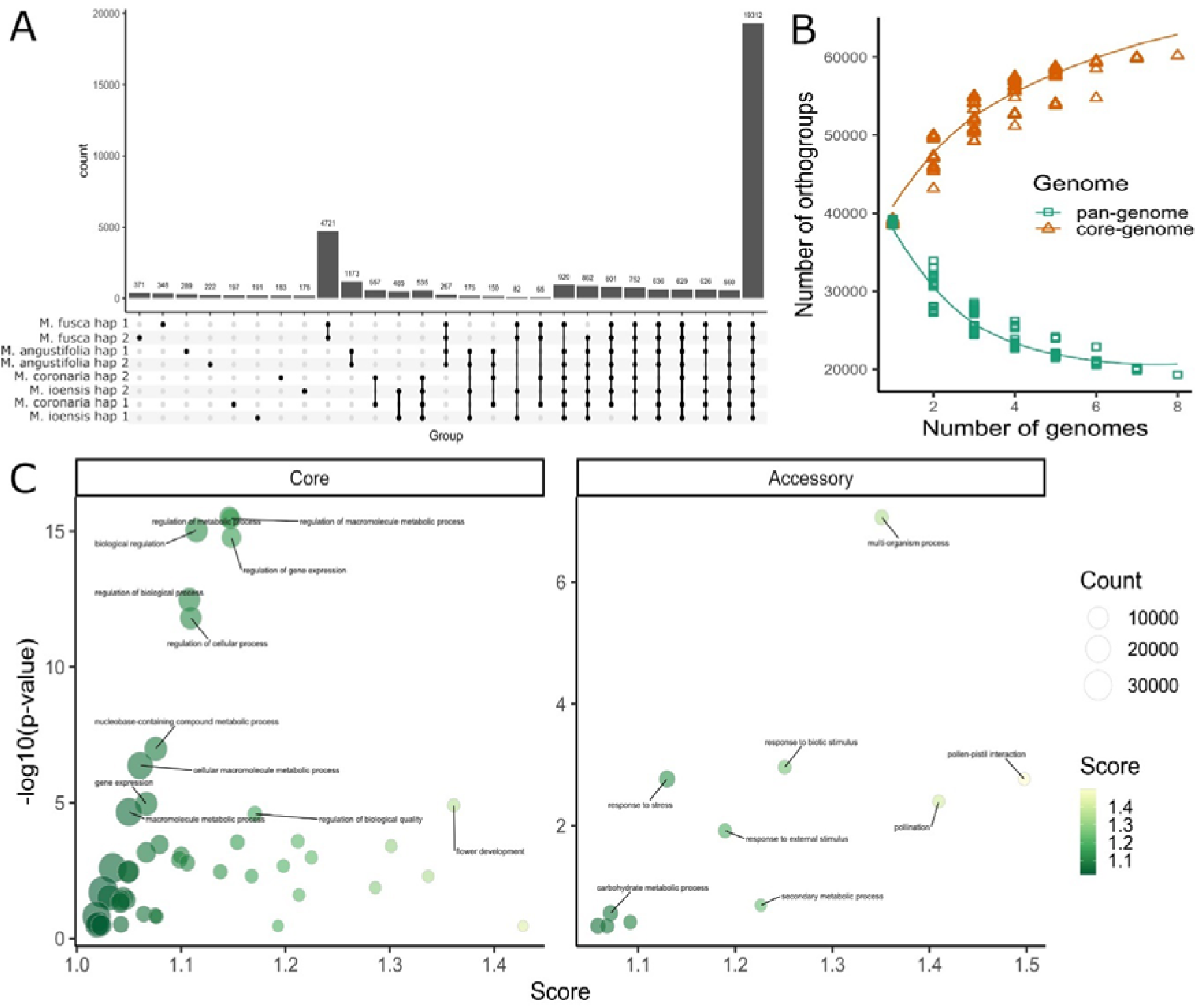
Pangenome of North American *Malus* species. **A)** Gene family based pangenome including eight haplomes of the four North American *Malus* species. **B)** Pan- and core genome models based on the orthogroups identified among the eight haplomes. **C)** Significantly enriched GO terms in orthogroups of the core and accessory genomes.

Orthogroups in the core genome encompass genes that showed enrichment of molecular mechanisms that are critical for the maintenance of basic metabolic functions, whereas genes in the accessory genome showed enrichment for mechanisms typically involved in adaptation to changing environments and stress. Genes involved in regulation of different metabolic process and gene expression showed most significant enrichment in core genome compared to the pangenome. Also, flower development, growth, and differentiation genes were significantly enriched in core genome and showed relatively high ratio between the count of the genes in core genome compared to those in pangenome. In contrast, genes involved in multi-organism processes, response to biotic and external stimulus, stress, as well as secondary metabolic processes, and pollination were enriched in the accessory genome. Finally, accession-specific genomes showed lower level or no enrichment, as only *M. ioensis*, *M.coronaria*, and *M. angustifolia* showed enrichment of genes involved in processes such as receptor activity, molecule binding, and signal transduction (Fig. S3).

Alignment of *M. coronaria*, *M. ioensis*, and *M. fusca* haplome 1 and 2 assemblies along the GDDH13 reference assembly [31] identified SVs and SV hotspots spanning intra- and intergenic genomic regions. Over 24,000 SVs identified between the two haplomes in each species resulted in a total length of 102 Mb, 103 Mb, and 85 Mb of SVs per *M. coronaria*, *M. ioensis*, and *M. fusca* genomes, respectively (Fig. 4 and Table S4). Insertions and deletions (InDels) were the most abundant, ranging from ∼9,000 to ∼14,000 of either insertions or deletions per haplome and represented up to 3.6 Mb per haplome (Fig. 4A). In contrast, inversions and translocations were less abundant with a minimum of 329 inversions in *M. ioensis* and a maximum of 373 in *M. fusca* that represented up to 87 Mb per genome. Between 4005 translocations in *M. fusca* and 4628 translocations in *M. coronaria* covered up to 13 Mb of the haplomes. An intersection between 400 kb windows along each of the assemblies and the SVs within these windows identified over 360 hotspots per genome containing up to 37 SVs per hospot (Fig. 4B). These hotspots overlapped with the regions with a higher number of genes (Pearson correlation coefficient *R*=0.53, *p* value<2.2E-16; Fig. S4), although significant functional enrichments of these genes were not found.

**Figure 4:**
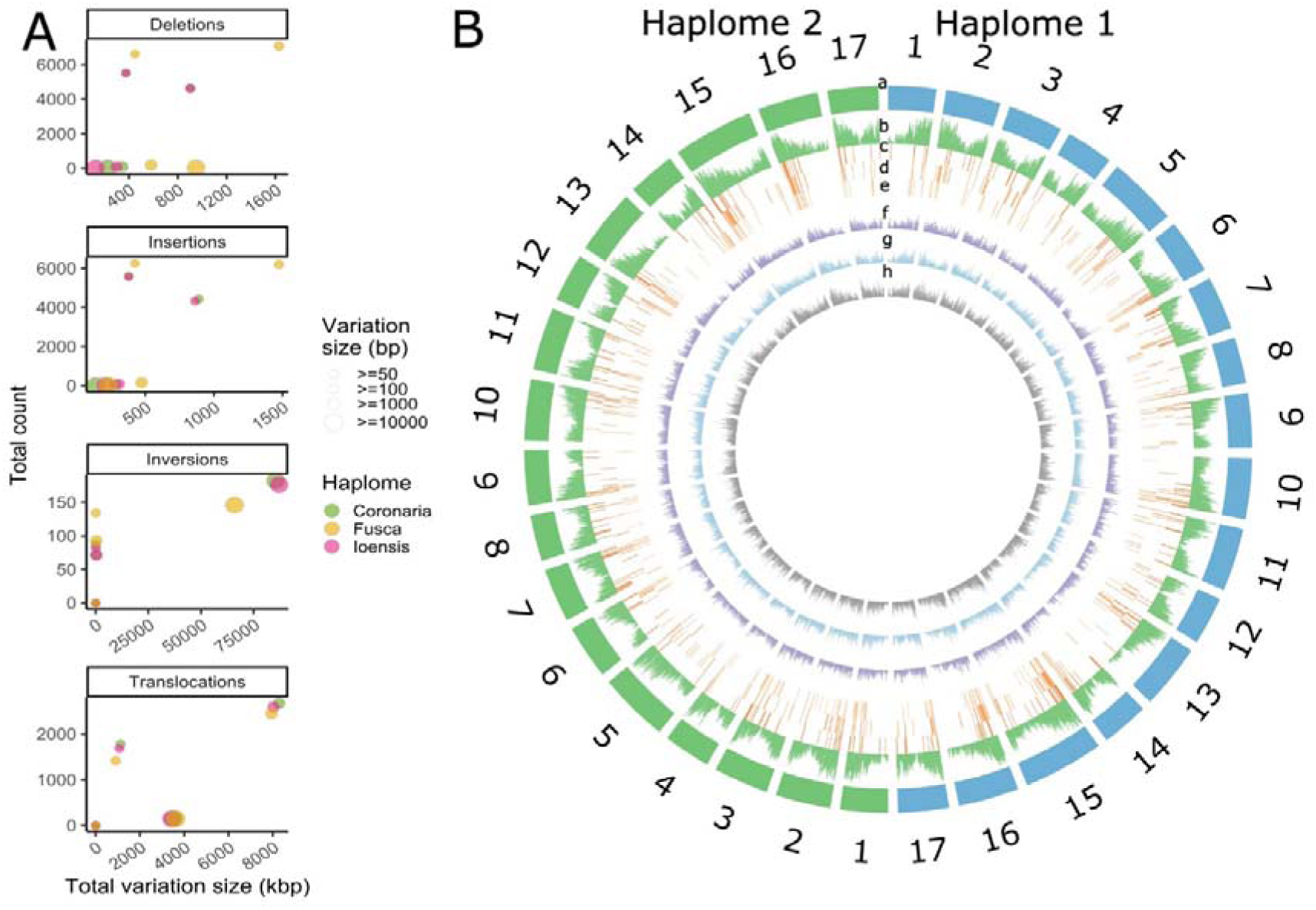
SVs in haplomes of *Malus coronaria* PI590014, *M. ioensis* PI590015, and *M. fusca*, compared to the reference GDDH13 genome. **A)** Total number, length, and distribution of deletions, insertions, inversions and translocations. **B)** SVs density along 400-bp windows. a, chromosomes. **b**, gene density, **c-e**, SV hotspots in haplomes of *M. coronaria* **(c),** *M. ioensis* (d) and *M. fusca* **(e)**. f-h, SVs in haplomes of *M. coronaria* **(f)**, *M. ioensis* **(g)**, and *M. fusca* **(h)**, compared to GDDH13.

## Discussion

This study presents high-quality phased chromosome-level genome assemblies of two North American *Malus* species and the first gene family based pangenome of North American *Malus* specie. The assembled *M. coronaria* and *M. ioensis* genomes had BUSCO completeness over 97.2%, k-mer completeness over 95.8%, and QV over 66.2. The quality of these genomes is comparable to the previously assembled genomes of *M. angustifolia* and *M. fusca*, which are native to North America as well [3,5], and other recent publications of high-quality *M. domestica* genomes and their close relatives [9,10,17,24,55]. The phased plant genomes enable studies of intragenomic variations, which could be linked with specific traits. For example, horticultural traits were associated to copy number variations and SVs of multiple *Malus* genomes [10], and fruit color and fruit acidity were associated with allele specific expression and allelic sequence variation in ‘Gala’, *M. sieversii*, and *M. sylvestris* apples [9]. Using the pangenome, we have identified gene functions that are core and accessory to the North American *Malus*, which can be studied further to understand the genetics underlying their specific traits such as distinct architecture, morphology, flowering patterns, and stress responses. Further dedicated phenotypic evaluation of *M. coronaria* and *M. ioensis* will enable such studies of North American *Malus* species. In addition, the addition of *M. coronaria* and *M. ioensis* genomes to the existing North American *Malus* assemblies offers opportunities to further explore their pangenomic landscape. This will aid further evolutionary studies, as well as development of novel markers for breeding purposes [8].

Phylogeny analysis of the *M. coronaria* and *M. ioensis* showed that these two accessions are highly related to *M. angustifolia* but differ from *M. fusca*. This fits their geographic origin, as *M. coronaria* and *M. ioensis* accessions were sampled from a close geographic proximity in central-eastern US [56], that was closer to south-eastern *M. angustifolia* than the north-western US sampling locations of *M. fusca* [3,5]. The three accessions from the east had a lower total number of macro-structural variations in genic regions among themselves compared with *M. fusca*, which could further suggest their lower relatedness with the western *M. fusca* species [3]. In contrast, close genetic relatedness, high level of macrosynteny, and the geographic proximity of *M. coronaria*, *M. ioensis*, and *M. angustifolia* indicates that the species from *Malus* section *Chloromeles* are conspecific [2]. Also, the large evolutionary distance from *M. domestica*, *M. sieversii*, and *M. sylvestris* further suggests that these species have limited cross-compatibility and geographic niche, and only rarely cross in nature [57].

Gene family based pangenome construction using protein sequences of two haplomes of *M. coronaria*, *M. ioensis*, *M. angustifolia*, and *M. fusca* identified core and accessory genes that describe a substantial diversity among North American *Malus* species, and suggested molecular functions critical for their evolution. Pangenome contained 60,211 orthogroups of 340,087 genes, and almost half of these genes were found as core to all the studied accessions and haplomes, whereas approximately a third of the genes were part of the accessory genome and less than 5% of genes were accession specific. Addition of more haplomes to the analysis only slightly decreased the core genome size, whereas the increase of the pangenome has been larger than the change in the core genome from seven to eight genomes. This indicates that core diversity was sufficiently described, whereas additional genomes should be added to further improve the representativeness of the pangenome. Core genome size fraction in relation to pangenome was equal to the previous study of thirteen *Malus* accessions [10], and similar to the core genome fraction in some other fruit crops including blueberry, cranberry [12] and strawberry [58], whereas it was lower than core genome fraction among ‘Gala’, *M. sieversii*, and *M. sylvestris* (81.3–87.3%) [9]. Core genes were enriched for basic cellular and metabolic processes and growth and development, indicating that basal metabolism and molecular functions are conserved across different *Malus* species [10]. In contrast, accessory genes were enriched for secondary metabolism, stress response, and interactions with other organisms, similarly to other *Malus* accessions [10] and *Vaccinium* [12]. These genes represent a gene pool that is important for species’ adjustment to different environments and might encompass alleles valuable for breeding for different agronomic traits such as their distinct morphology, flowering pattern, disease resistance and abiotic stress tolerance [10,12]. Finally, species-specific genes showed only a low level of enriched genes that further pinpoint towards high relatedness of the studied species, although further analysis of gene speciation will be required by incorporating a larger intraspecies diversity.

The identified SVs were concentrated in the regions of the genome that have a high gene density and could be associated with variation in specific traits [59]. Some of the North American *Malus* are reputed for their different morphology, delayed flowering, disease resistance, climate adaptation, and the ability to grow on sandy waterlogged soils [4,5,57]. The identified SVs can serve as the foundation of markers for the development of a genotyping platform that can be used in the future breeding efforts to utilize North American species for apple cultivar development [1]. Our study demonstrated that InDels were the most abundant among the identified SVs, whereas translocations and inversions represented the largest portion of the genome. These SVs could be utilized for further studies of specific traits, as in the past, SVs were demonstrated to be involved in fruit color definition, as well as they are associated with fruit quality and (a)biotic stress tolerance [8,10]. Other SVs might be impactful as well, as resistance to fire blight in *M. fusca* was suggested to be associated with copy number variation of *R* genes on linkage group 10 [5]. Further studies are necessary to identify the most important SVs in the genomes of North American *Malus* species.

Native North-American *Malus* species historically represent an important nourishment food source for the indigenous inhabitants of the New World [60]. *M. fusca* in the west, and *M. coronaria*, *M. ioensis*, and *M. angustifolia* in the east were not propagated by grafting or cultivated in orchards, but these crab apples are still particularly useful for roasting, or as preserved in maple syrup, or boiled into sauce [60]. Today, the genetic information for all four native apples is accessible as we have constructed high-quality chromosome-level phased genome assemblies of *M. coronaria* and *M. ioensis* and the first gene family based pangenome of North American *Malus* species. This resulted in further establishment of evolutionary relationships among *Malus* species of North America and with other *Malus* species, and the identification of SVs. The high quality of the two new genome assemblies offers opportunities to further utilize them in dedicated *Malus* inter- and intraspecific pangenomic studies. Individual assemblies, as well as the gene family based pangenome could be instrumental in the future to obtain a deeper understanding of plant evolution and adaptation, and to identify genes and SVs for apple breeding.

### Data availability statement

The datasets supporting the conclusions of this article comprising HiFi and Hi-C raw reads and haplome assemblies of *M. coronaria* and *M. ioensis* has been deposited in the NCBI under the Bioproject accession numbers PRJNA1098636, PRJNA1098637, PRJNA1098638, and PRJNA1098639 and will be accessible upon publication of the article. The datasets used in this article will also be made available in the Genomic Database for Rosaceae (GDR) after acceptance of the manuscript.

## Acknowledgements

We gratefully acknowledge the stewardship of Indigenous nations over North American *Malus* native species and genetic diversity, honoring their profound connection to the land and its inhabitants. This research was supported by the USDA-NIFA Specialty Crop Research Initiative (SCRI) grant 1023572 (subaward RC111414A), and the New York State Specialty Crop Block Grant “SCG 18 008”. During the duration of this research, Anže Švara was supported by BAEF (Belgian American Educational Foundation) as Ernest du Bois Fellow. We acknowledge the help of Della Cobb-Smith during sample collection for DNA extraction and sequencing, and IT support of Dr. Qi Sun during data analysis.

## Declarations

Not applicable

## Conflicts of Interest

The authors declare that they have no competing interests.

## Author Contributions

A.K. designed the experiment and supervised the research. A.S performed genome assemblies and pangenome analysis. H.S. performed genome annotation. A.S. wrote the manuscript. A.K, and Z.F. revised the manuscript. All authors have read and approved the manuscript.

## Supplementary Files

**Figure S1:**
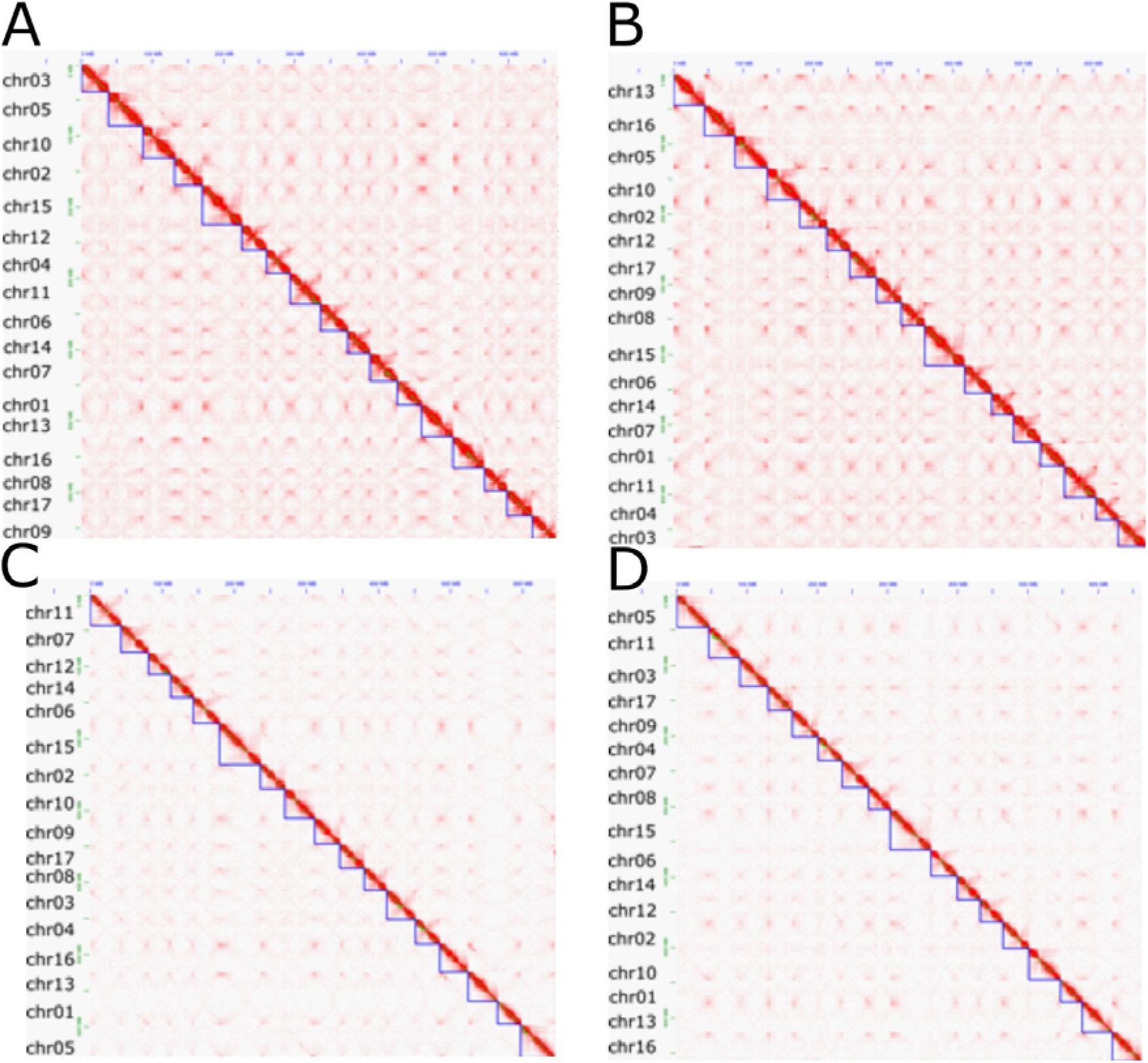
Chromatin contact maps of *Malus coronaria* accession PI590014 A) haplome 1 and B) 2, and *Malus ioensis* accession PI590015 haplome C) 1 and D) 2.

**Figure S2:**
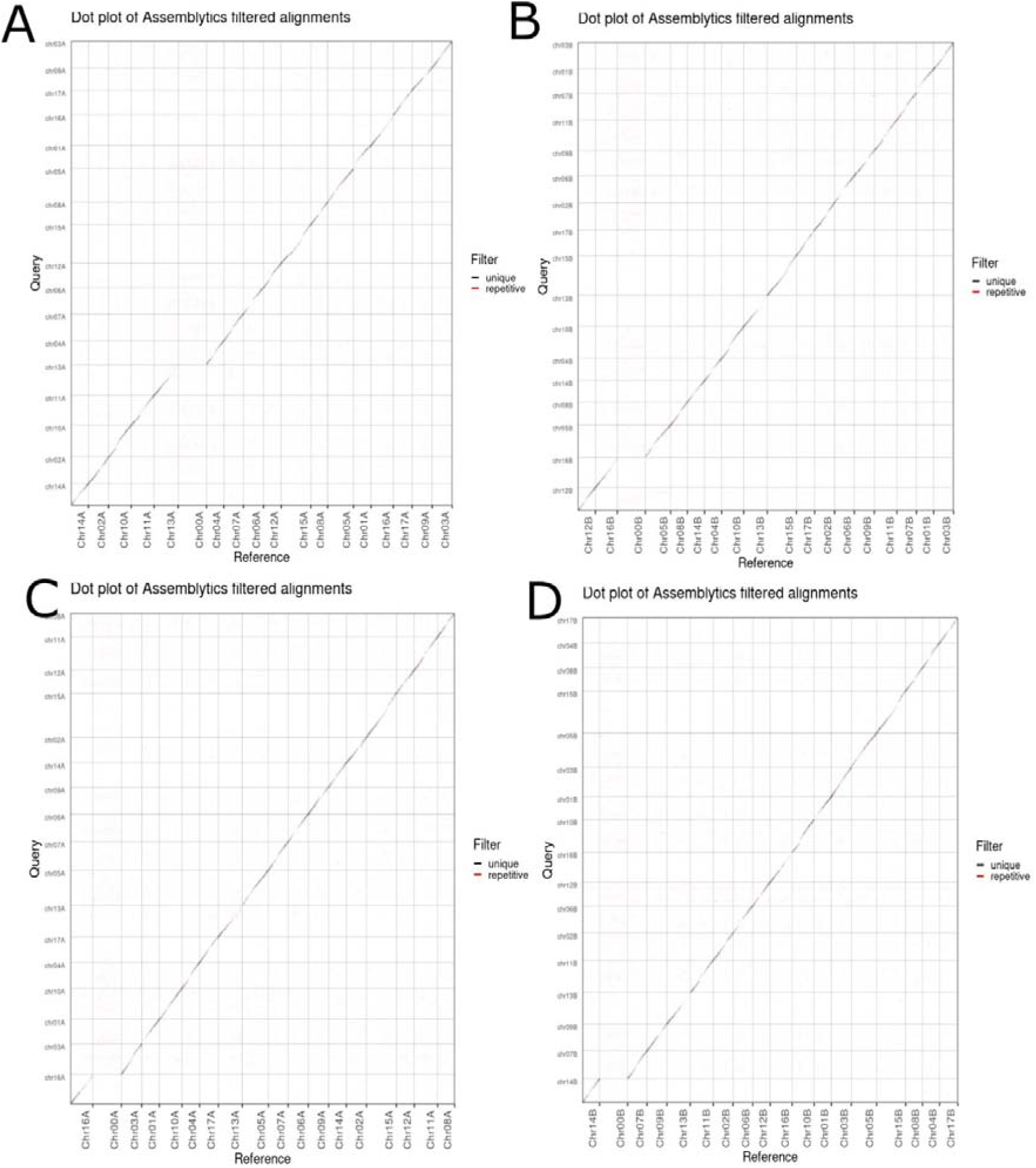
Collinearity between *Malus coronaria* accession PI590014 haplomes A) 1 and B) 2 and C) haplome 1 and D) 2 of *Malus ioensis* accession PI590015 with GDDH13 genome [31].

**Fig S3:**
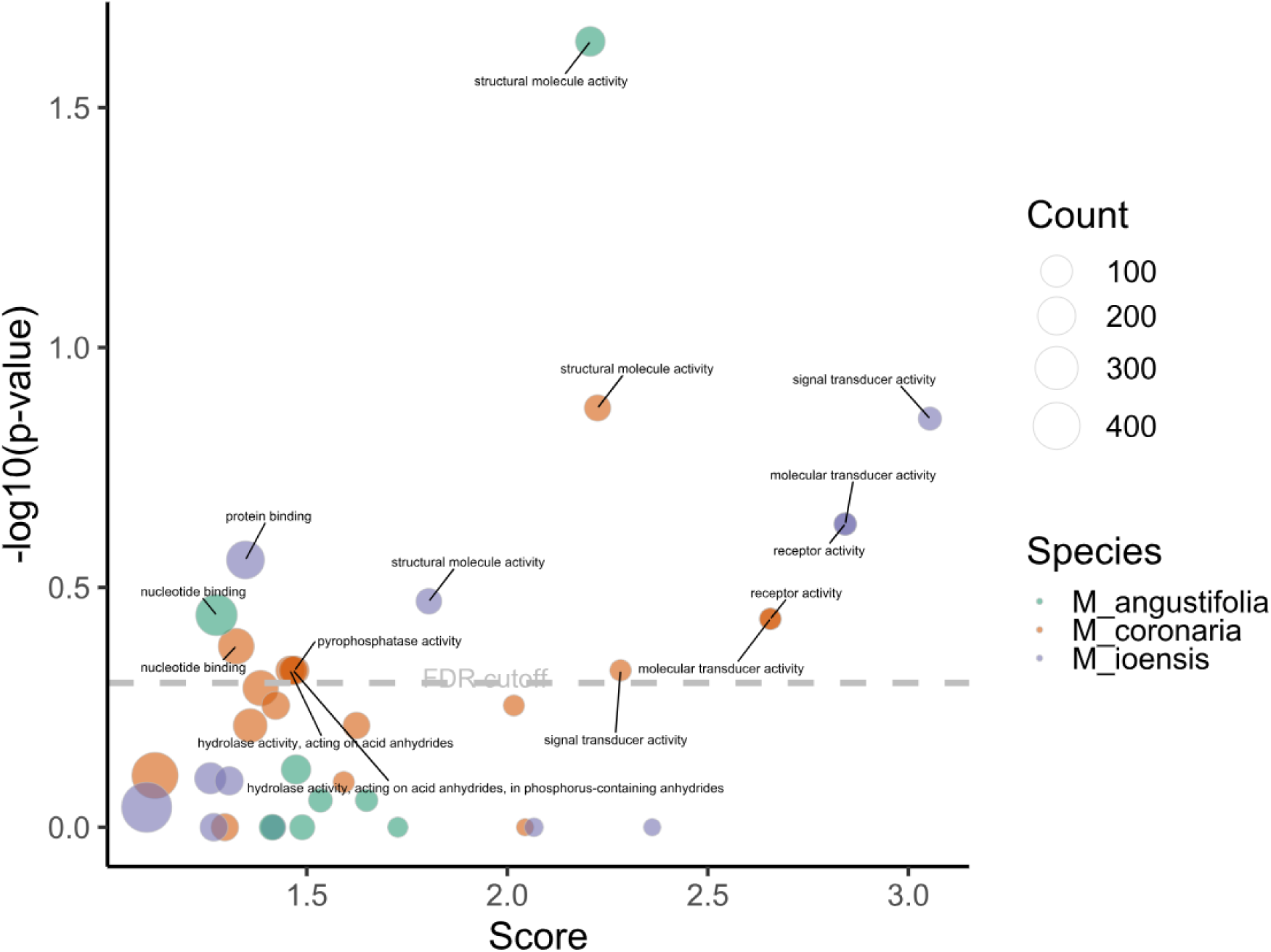
Enrichment of processes associated with genes of the orthogroups belonging to species-specific genomes of *Malus coronaria*, *M. ioensis*, *M. angustifolia*, and *M. fusca* (no significant enrichment found).

**Fig S4:**
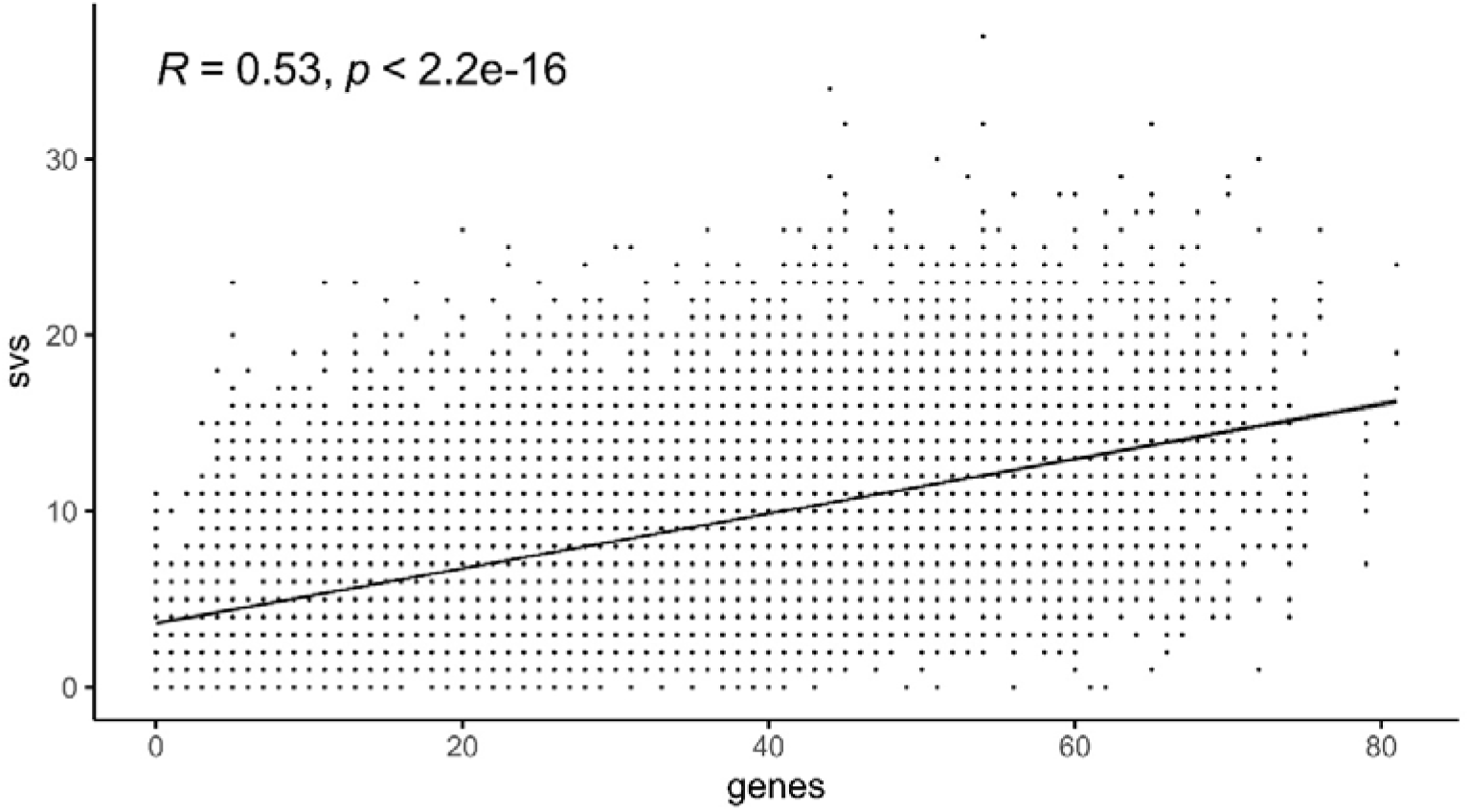
Correlation plot showing the number of genes and structural variations within 400 kb regions along the *Malus coronaria*, *M. ioensis*, and *M. fusca* genomes.

**Table S1:**
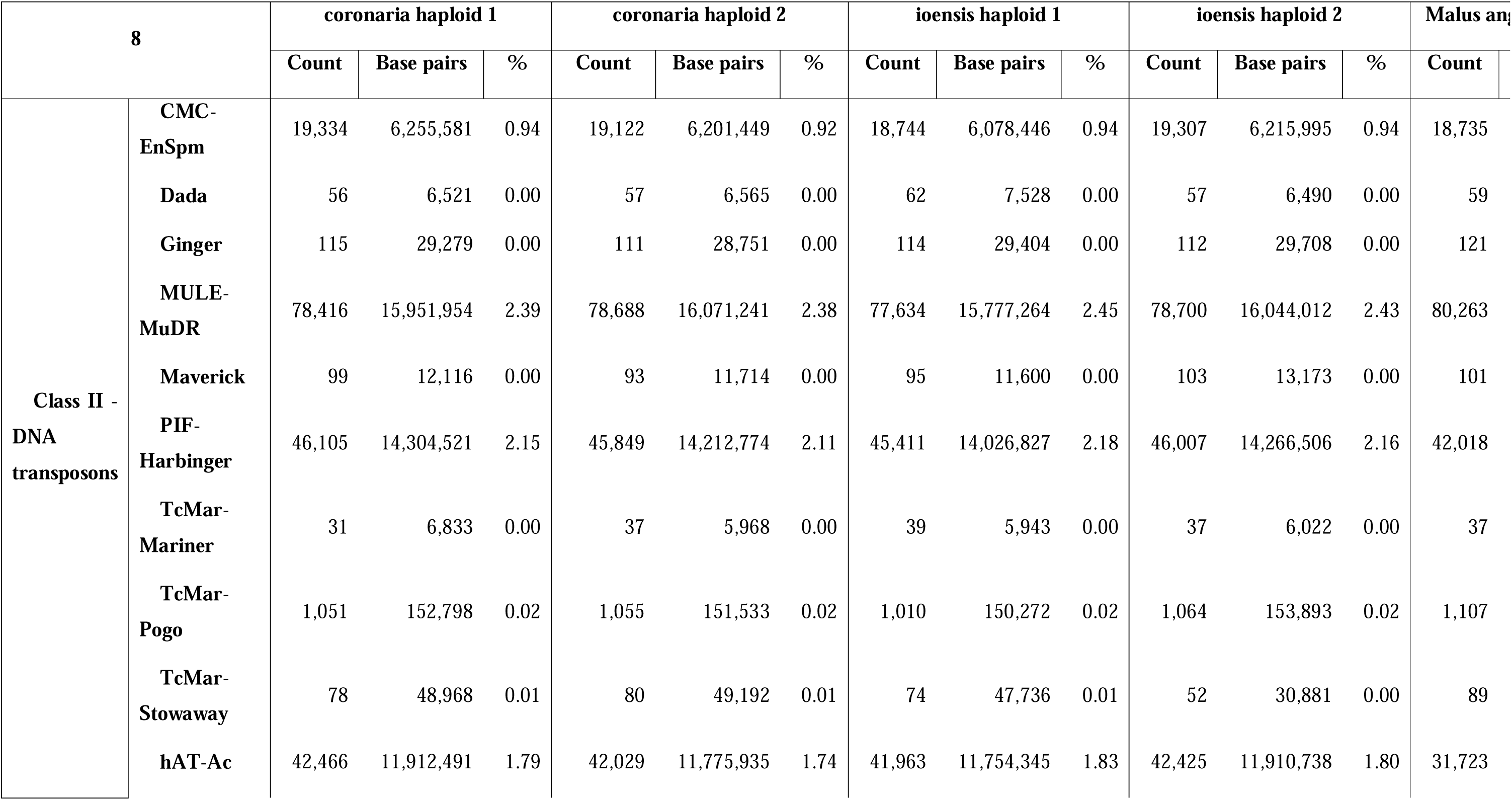
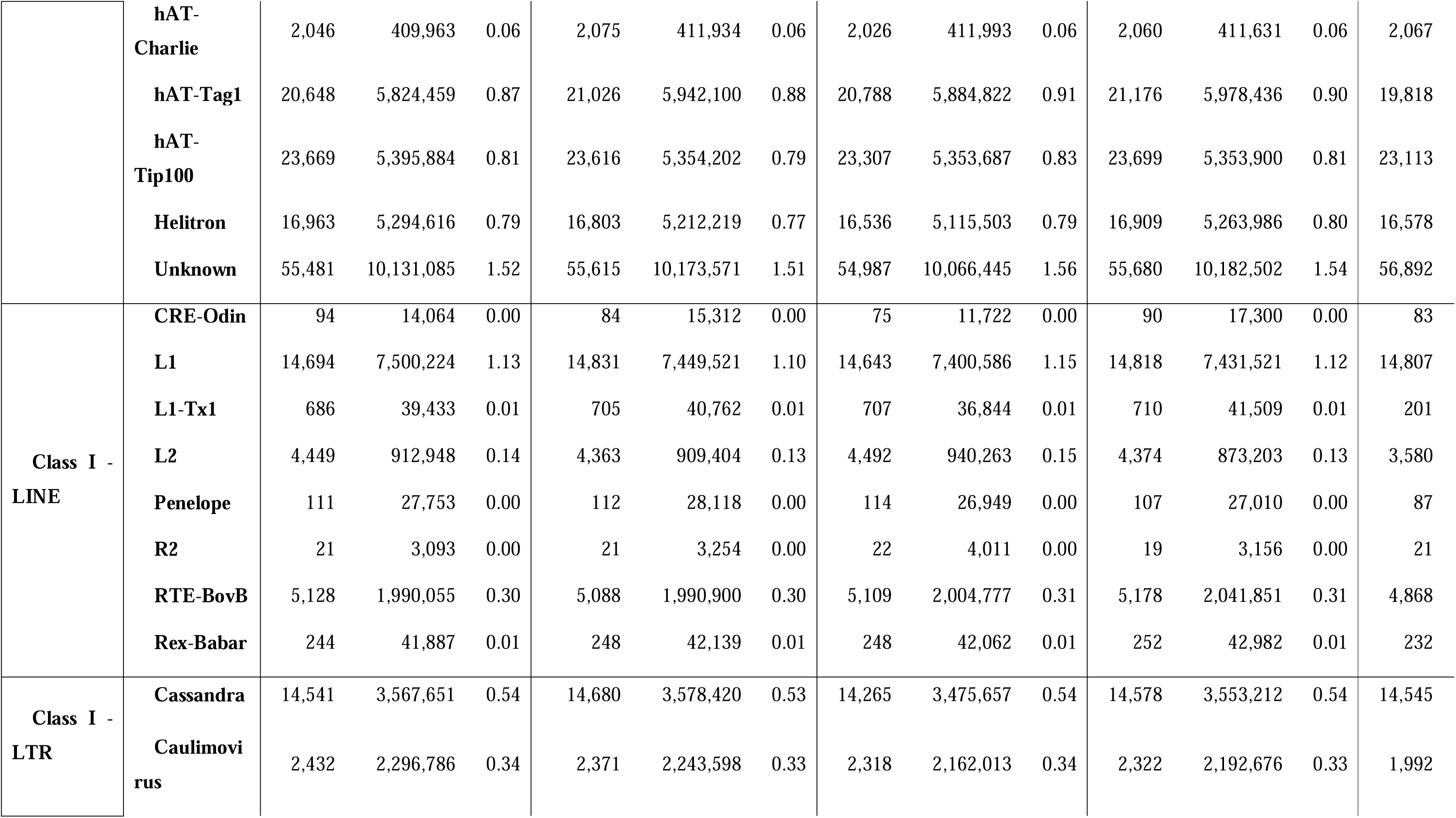
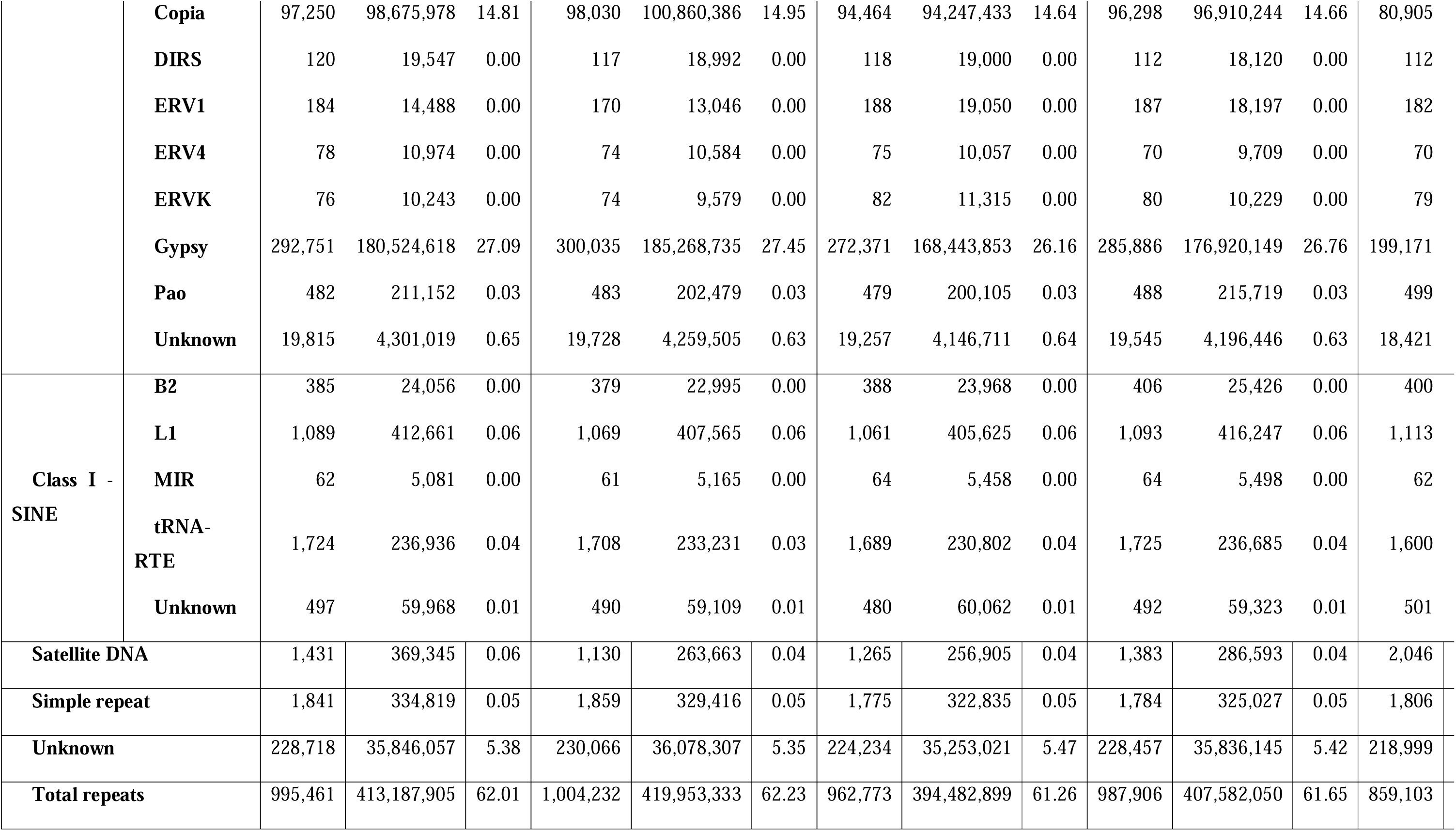
Repeats summary of *Malus coronaria* (PI590014) and *M. ioensis* (PI590015) haplomes, and of previously-assembled *M. angustifolia* and *M. fusca* haplomes [3,5].

| 8 |  | coronaria haploid 1 |  |  | coronaria haploid 2 |  |  | ioensis haploid 1 |  |  | ioensis haploid 2 |  |  | Malus angustifolia |
| --- | --- | --- | --- | --- | --- | --- | --- | --- | --- | --- | --- | --- | --- | --- |
|  |  | Count | Base pairs | % | Count | Base pairs | % | Count | Base pairs | % | Count | Base pairs | % | Count |
| Class II -<br>DNA<br>transposons | CMC-EnSpm | 19,334 | 6,255,581 | 0.94 | 19,122 | 6,201,449 | 0.92 | 18,744 | 6,078,446 | 0.94 | 19,307 | 6,215,995 | 0.94 | 18,735 |
|  | Dada | 56 | 6,521 | 0.00 | 57 | 6,565 | 0.00 | 62 | 7,528 | 0.00 | 57 | 6,490 | 0.00 | 59 |
|  | Ginger | 115 | 29,279 | 0.00 | 111 | 28,751 | 0.00 | 114 | 29,404 | 0.00 | 112 | 29,708 | 0.00 | 121 |
|  | MULE-MuDR | 78,416 | 15,951,954 | 2.39 | 78,688 | 16,071,241 | 2.38 | 77,634 | 15,777,264 | 2.45 | 78,700 | 16,044,012 | 2.43 | 80,263 |
|  | Maverick | 99 | 12,116 | 0.00 | 93 | 11,714 | 0.00 | 95 | 11,600 | 0.00 | 103 | 13,173 | 0.00 | 101 |
|  | PIF-Harbinger | 46,105 | 14,304,521 | 2.15 | 45,849 | 14,212,774 | 2.11 | 45,411 | 14,026,827 | 2.18 | 46,007 | 14,266,506 | 2.16 | 42,018 |
|  | TcMar-Mariner | 31 | 6,833 | 0.00 | 37 | 5,968 | 0.00 | 39 | 5,943 | 0.00 | 37 | 6,022 | 0.00 | 37 |
|  | TcMar-Pogo | 1,051 | 152,798 | 0.02 | 1,055 | 151,533 | 0.02 | 1,010 | 150,272 | 0.02 | 1,064 | 153,893 | 0.02 | 1,107 |
|  | TcMar-Stowaway | 78 | 48,968 | 0.01 | 80 | 49,192 | 0.01 | 74 | 47,736 | 0.01 | 52 | 30,881 | 0.00 | 89 |
|  | hAT-Ac | 42,466 | 11,912,491 | 1.79 | 42,029 | 11,775,935 | 1.74 | 41,963 | 11,754,345 | 1.83 | 42,425 | 11,910,738 | 1.80 | 31,723 |
|  | **hAT-Charlie** | 2,046 | 409,963 | 0.06 | 2,075 | 411,934 | 0.06 | 2,026 | 411,993 | 0.06 | 2,060 | 411,631 | 0.06 | 2,067 |
|  | **hAT-Tag1** | 20,648 | 5,824,459 | 0.87 | 21,026 | 5,942,100 | 0.88 | 20,788 | 5,884,822 | 0.91 | 21,176 | 5,978,436 | 0.90 | 19,818 |
|  | **hAT-Tip100** | 23,669 | 5,395,884 | 0.81 | 23,616 | 5,354,202 | 0.79 | 23,307 | 5,353,687 | 0.83 | 23,699 | 5,353,900 | 0.81 | 23,113 |
|  | **Helitron** | 16,963 | 5,294,616 | 0.79 | 16,803 | 5,212,219 | 0.77 | 16,536 | 5,115,503 | 0.79 | 16,909 | 5,263,986 | 0.80 | 16,578 |
|  | **Unknown** | 55,481 | 10,131,085 | 1.52 | 55,615 | 10,173,571 | 1.51 | 54,987 | 10,066,445 | 1.56 | 55,680 | 10,182,502 | 1.54 | 56,892 |
| **Class I - LINE** | **CRE-Odin** | 94 | 14,064 | 0.00 | 84 | 15,312 | 0.00 | 75 | 11,722 | 0.00 | 90 | 17,300 | 0.00 | 83 |
|  | **L1** | 14,694 | 7,500,224 | 1.13 | 14,831 | 7,449,521 | 1.10 | 14,643 | 7,400,586 | 1.15 | 14,818 | 7,431,521 | 1.12 | 14,807 |
|  | **L1-Tx1** | 686 | 39,433 | 0.01 | 705 | 40,762 | 0.01 | 707 | 36,844 | 0.01 | 710 | 41,509 | 0.01 | 201 |
|  | **L2** | 4,449 | 912,948 | 0.14 | 4,363 | 909,404 | 0.13 | 4,492 | 940,263 | 0.15 | 4,374 | 873,203 | 0.13 | 3,580 |
|  | **Penelope** | 111 | 27,753 | 0.00 | 112 | 28,118 | 0.00 | 114 | 26,949 | 0.00 | 107 | 27,010 | 0.00 | 87 |
|  | **R2** | 21 | 3,093 | 0.00 | 21 | 3,254 | 0.00 | 22 | 4,011 | 0.00 | 19 | 3,156 | 0.00 | 21 |
|  | **RTE-BovB** | 5,128 | 1,990,055 | 0.30 | 5,088 | 1,990,900 | 0.30 | 5,109 | 2,004,777 | 0.31 | 5,178 | 2,041,851 | 0.31 | 4,868 |
|  | **Rex-Babar** | 244 | 41,887 | 0.01 | 248 | 42,139 | 0.01 | 248 | 42,062 | 0.01 | 252 | 42,982 | 0.01 | 232 |
| **Class I - LTR** | **Cassandra** | 14,541 | 3,567,651 | 0.54 | 14,680 | 3,578,420 | 0.53 | 14,265 | 3,475,657 | 0.54 | 14,578 | 3,553,212 | 0.54 | 14,545 |
|  | **Caulimovirus** | 2,432 | 2,296,786 | 0.34 | 2,371 | 2,243,598 | 0.33 | 2,318 | 2,162,013 | 0.34 | 2,322 | 2,192,676 | 0.33 | 1,992 |
|  | **Copia** | 97,250 | 98,675,978 | 14.81 | 98,030 | 100,860,386 | 14.95 | 94,464 | 94,247,433 | 14.64 | 96,298 | 96,910,244 | 14.66 | 80,905 |
|  | **DIRS** | 120 | 19,547 | 0.00 | 117 | 18,992 | 0.00 | 118 | 19,000 | 0.00 | 112 | 18,120 | 0.00 | 112 |
|  | **ERV1** | 184 | 14,488 | 0.00 | 170 | 13,046 | 0.00 | 188 | 19,050 | 0.00 | 187 | 18,197 | 0.00 | 182 |
|  | **ERV4** | 78 | 10,974 | 0.00 | 74 | 10,584 | 0.00 | 75 | 10,057 | 0.00 | 70 | 9,709 | 0.00 | 70 |
|  | **ERVK** | 76 | 10,243 | 0.00 | 74 | 9,579 | 0.00 | 82 | 11,315 | 0.00 | 80 | 10,229 | 0.00 | 79 |
|  | **Gypsy** | 292,751 | 180,524,618 | 27.09 | 300,035 | 185,268,735 | 27.45 | 272,371 | 168,443,853 | 26.16 | 285,886 | 176,920,149 | 26.76 | 199,171 |
|  | **Pao** | 482 | 211,152 | 0.03 | 483 | 202,479 | 0.03 | 479 | 200,105 | 0.03 | 488 | 215,719 | 0.03 | 499 |
|  | **Unknown** | 19,815 | 4,301,019 | 0.65 | 19,728 | 4,259,505 | 0.63 | 19,257 | 4,146,711 | 0.64 | 19,545 | 4,196,446 | 0.63 | 18,421 |
| **Class I - SINE** | **B2** | 385 | 24,056 | 0.00 | 379 | 22,995 | 0.00 | 388 | 23,968 | 0.00 | 406 | 25,426 | 0.00 | 400 |
|  | **L1** | 1,089 | 412,661 | 0.06 | 1,069 | 407,565 | 0.06 | 1,061 | 405,625 | 0.06 | 1,093 | 416,247 | 0.06 | 1,113 |
|  | **MIR** | 62 | 5,081 | 0.00 | 61 | 5,165 | 0.00 | 64 | 5,458 | 0.00 | 64 | 5,498 | 0.00 | 62 |
|  | **tRNA-RTE** | 1,724 | 236,936 | 0.04 | 1,708 | 233,231 | 0.03 | 1,689 | 230,802 | 0.04 | 1,725 | 236,685 | 0.04 | 1,600 |
|  | **Unknown** | 497 | 59,968 | 0.01 | 490 | 59,109 | 0.01 | 480 | 60,062 | 0.01 | 492 | 59,323 | 0.01 | 501 |
| **Satellite DNA** |  | 1,431 | 369,345 | 0.06 | 1,130 | 263,663 | 0.04 | 1,265 | 256,905 | 0.04 | 1,383 | 286,593 | 0.04 | 2,046 |
| **Simple repeat** |  | 1,841 | 334,819 | 0.05 | 1,859 | 329,416 | 0.05 | 1,775 | 322,835 | 0.05 | 1,784 | 325,027 | 0.05 | 1,806 |
| **Unknown** |  | 228,718 | 35,846,057 | 5.38 | 230,066 | 36,078,307 | 5.35 | 224,234 | 35,253,021 | 5.47 | 228,457 | 35,836,145 | 5.42 | 218,999 |
| **Total repeats** |  | 995,461 | 413,187,905 | 62.01 | 1,004,232 | 419,953,333 | 62.23 | 962,773 | 394,482,899 | 61.26 | 987,906 | 407,582,050 | 61.65 | 859,103 |

**Table S2:** N repeats summary of *Malus coronaria* (PI590014) and *M. ioensis* (PI590015) haplomes, and of previously-assembled *M. angustifolia* and *M. fusca* haplomes [3,5].

| Genome | Assembly size | N gap total length |
| --- | --- | --- |
| coronaria haploid 1 | 666,594,626 | 304,500 |
| coronaria haploid 2 | 675,532,332 | 295,000 |
| ioensis haploid 1 | 644,188,171 | 288,500 |
| ioensis haploid 2 | 661,368,478 | 294,500 |
| Malus angustifolia haploid 1 | 705,089,958 | 234,793,148 |
| Malus angustifolia haploid 2 | 703,144,148 | 234,311,768 |
| Malus fusca haploid 1 | 637,459,684 | 3,200 |
| Malus fusca haploid 2 | 631,922,950 | 2,800 |

**Table S3:** Gene annotation statistics of.

| BUSCO estimate | coronaria haploid 1 |  | coronaria haploid 2 |  | coronaria phased diploid |  | ioensis haploid 1 |  | ioensis haploid 2 |  | ioensis phased diploid |  | Malus angustifolia haploid 1 |  | Malus angustifolia haploid 2 |  | Malus angustifolia phased diploid |  | Malus fusca haploid 1 |  | Malus fusca haploid 2 |  | Malus fusca phased diploid |  |
| --- | --- | --- | --- | --- | --- | --- | --- | --- | --- | --- | --- | --- | --- | --- | --- | --- | --- | --- | --- | --- | --- | --- | --- | --- |
| Data set | genome | genes | genome | genes | genome | genes | genome | genes | genome | genes | genome | genes | genome | genes | genome | genes | genome | genes | genome | genes | genome | genes | genome | genes |
| All complete % | 97.77 | 96.59 | 97.71 | 96.90 | 98.39 | 98.02 | 97.15 | 95.79 | 97.89 | 97.09 | 98.51 | 97.89 | 98.70 | 97.71 | 98.57 | 97.77 | 98.95 | 98.45 | 98.70 | 98.02 | 98.82 | 98.33 | 98.82 | 98.64 |
| **Single**<br>**compl**<br>**ete %** | 62.<br>45 | 65<br>.1<br>8 | 62.<br>76 | 65<br>.3<br>0 | 2.4<br>8 | 3.<br>47 | 61.<br>96 | 64<br>.7<br>5 | 62.<br>14 | 64<br>.3<br>1 | 3.2<br>2 | 3.<br>66 | 62.<br>02 | 64<br>.7<br>5 | 62.<br>14 | 64<br>.5<br>6 | 1.5<br>5 | 2.<br>54 | 61.<br>65 | 63<br>.1<br>4 | 61.<br>52 | 63<br>.2<br>6 | 0.8<br>1 | 1.92 |
| **Multi**<br>**ple**<br>**compl**<br>**ete %** | 35.<br>32 | 31<br>.4<br>2 | 34.<br>95 | 31<br>.6<br>0 | 95.<br>91 | 94<br>.5<br>4 | 35.<br>19 | 31<br>.0<br>5 | 35.<br>75 | 32<br>.7<br>7 | 95.<br>29 | 94<br>.2<br>3 | 36.<br>68 | 32<br>.9<br>7 | 36.<br>44 | 33<br>.2<br>1 | 97.<br>40 | 95<br>.9<br>1 | 37.<br>05 | 34<br>.8<br>8 | 37.<br>30 | 35<br>.0<br>7 | 98.<br>01 | 96.72 |
| **Frag**<br>**mente**<br>**d %** | 1.0<br>5 | 0.<br>93 | 0.9<br>9 | 0.<br>74 | 0.8<br>1 | 0.<br>43 | 1.1<br>2 | 0.<br>99 | 0.8<br>7 | 0.<br>37 | 0.8<br>1 | 0.<br>37 | 0.5<br>6 | 0.<br>56 | 0.6<br>2 | 0.<br>25 | 0.5<br>6 | 0.<br>37 | 0.6<br>8 | 0.<br>56 | 0.6<br>8 | 0.<br>50 | 0.6<br>8 | 0.43 |
| **Missi**<br>**ng %** | 1.1<br>8 | 2.<br>48 | 1.3<br>0 | 2.<br>35 | 0.8<br>1 | 1.<br>55 | 1.7<br>3 | 3.<br>22 | 1.2<br>4 | 2.<br>54 | 0.6<br>8 | 1.<br>73 | 0.7<br>4 | 1.<br>73 | 0.8<br>1 | 1.<br>98 | 0.5<br>0 | 1.<br>18 | 0.6<br>2 | 1.<br>43 | 0.5<br>0 | 1.<br>18 | 0.5<br>0 | 0.93 |
| **BUSC**<br>**O**<br>**gene**<br>**numb**<br>**er** | 161<br>4 | 16<br>14 | 161<br>4 | 16<br>14 | 161<br>4 | 16<br>14 | 161<br>4 | 16<br>14 | 161<br>4 | 16<br>14 | 161<br>4 | 16<br>14 | 161<br>4 | 16<br>14 | 161<br>4 | 16<br>14 | 161<br>4 | 16<br>14 | 161<br>4 | 16<br>14 | 161<br>4 | 16<br>14 | 161<br>4 | 1614 |
| **All**<br>**compl**<br>**ete** | 157<br>8 | 15<br>59 | 157<br>7 | 15<br>64 | 158<br>8 | 15<br>82 | 156<br>8 | 15<br>46 | 158<br>0 | 15<br>67 | 159<br>0 | 15<br>80 | 159<br>3 | 15<br>77 | 159<br>1 | 15<br>78 | 159<br>7 | 15<br>89 | 159<br>3 | 15<br>82 | 159<br>5 | 15<br>87 | 159<br>5 | 1592 |
| **Single**<br>**compl** | 100<br>8 | 10<br>52 | 101<br>3 | 10<br>54 | 40 | 56 | 100<br>0 | 10<br>45 | 100<br>3 | 10<br>38 | 52 | 59 | 100<br>1 | 10<br>45 | 100<br>3 | 10<br>42 | 25 | 41 | 995 | 10<br>19 | 993 | 10<br>21 | 13 | 31 |
| **ete** |  |  |  |  |  |  |  |  |  |  |  |  |  |  |  |  |  |  |  |  |  |  |  |  |
| **Multi**<br>**ple**<br>**compl**<br>**ete** | 570 | 50<br>7 | 564 | 51<br>0 | 154<br>8 | 15<br>26 | 568 | 50<br>1 | 577 | 52<br>9 | 153<br>8 | 15<br>21 | 592 | 53<br>2 | 588 | 53<br>6 | 157<br>2 | 15<br>48 | 598 | 56<br>3 | 602 | 56<br>6 | 158<br>2 | 1561 |
| **Frag**<br>**mente**<br>**d** | 17 | 15 | 16 | 12 | 13 | 7 | 18 | 16 | 14 | 6 | 13 | 6 | 9 | 9 | 10 | 4 | 9 | 6 | 11 | 9 | 11 | 8 | 11 | 7 |
| **Missi**<br>**ng** | 19 | 40 | 21 | 38 | 13 | 25 | 28 | 52 | 20 | 41 | 11 | 28 | 12 | 28 | 13 | 32 | 8 | 19 | 10 | 23 | 8 | 19 | 8 | 15 |

**Table S4:**
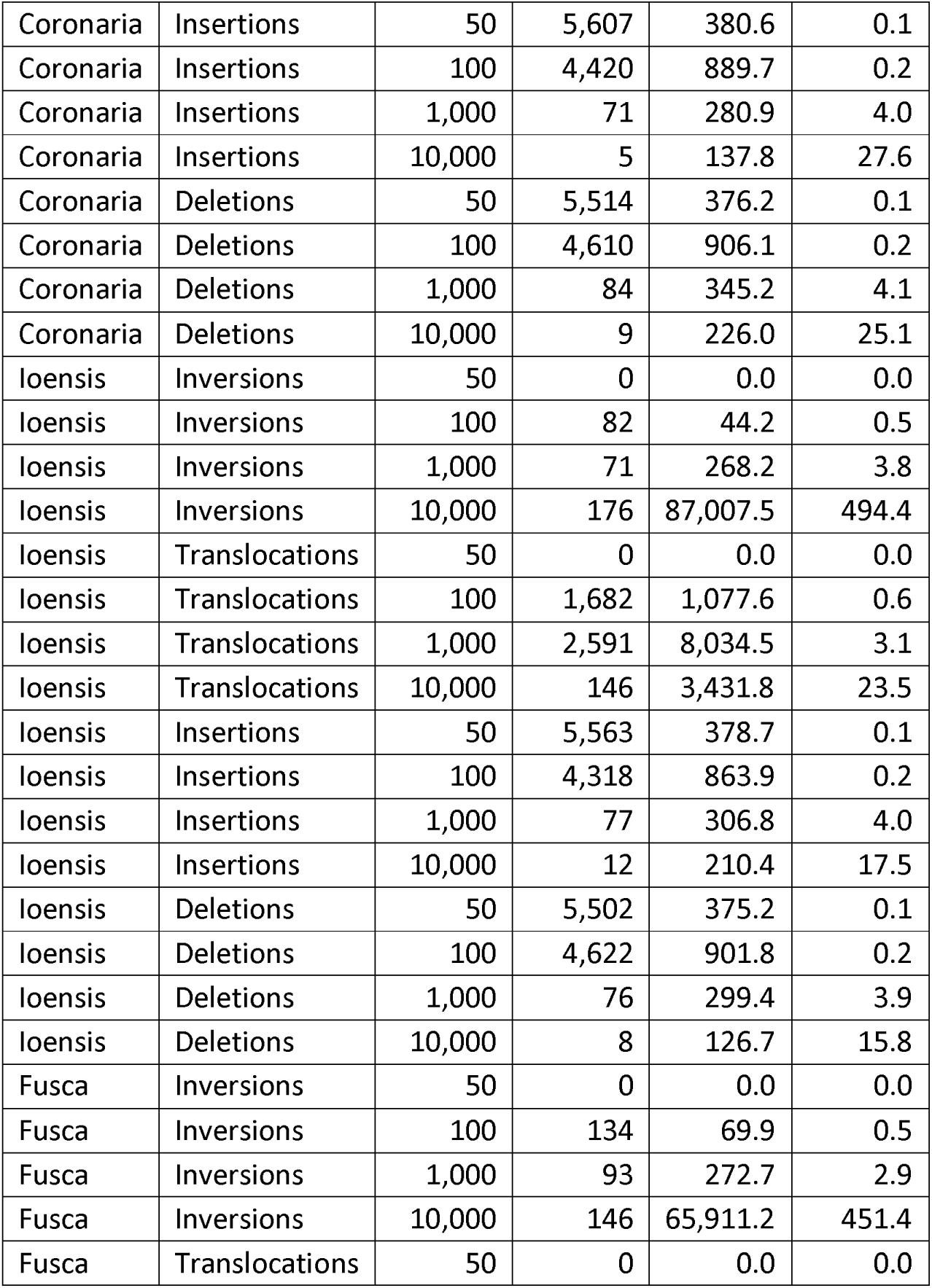
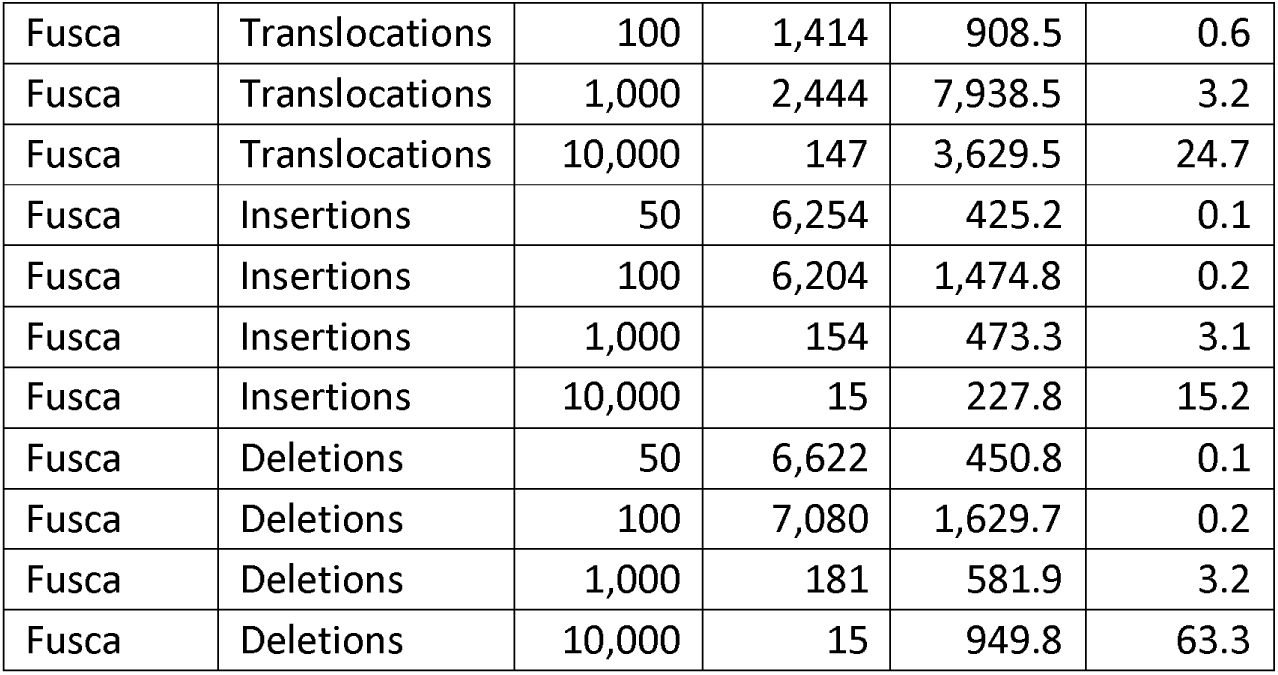
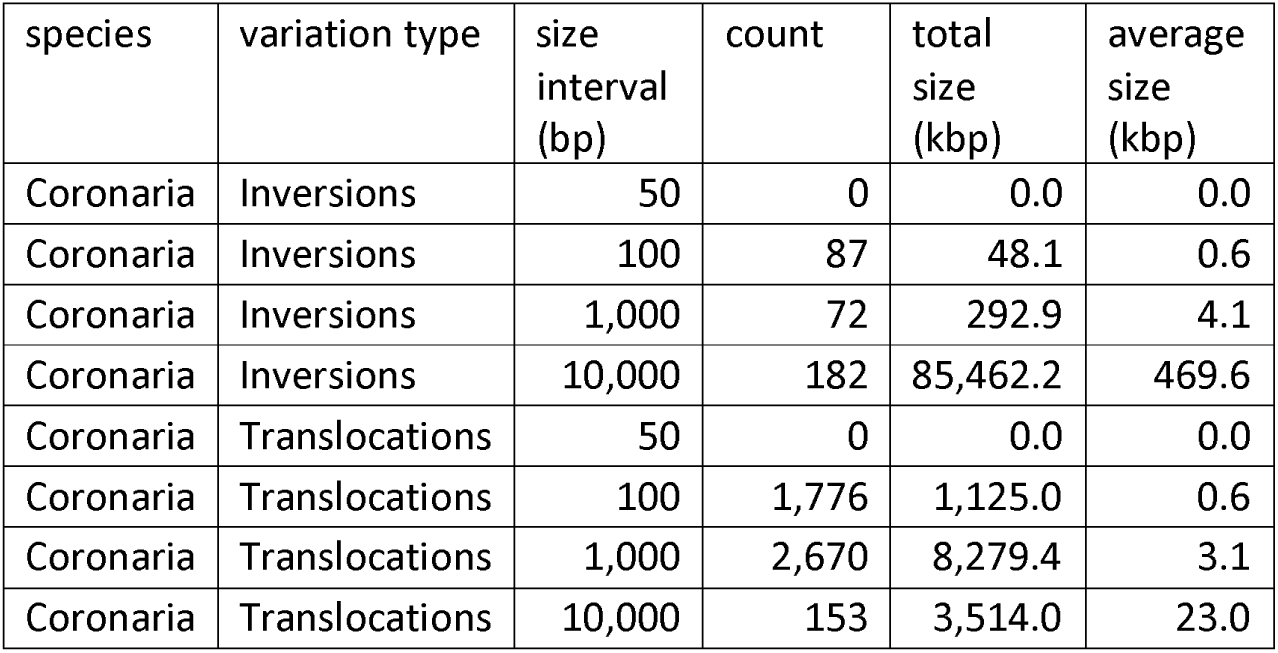
Summary statistics of the identified structural variation in *Malus coronaria* PI590014, *M. ioensis* PI590015, and *M. fusca* haplomes 1 and 2.

## Notes

### Competing Interest Statement

The authors have declared no competing interest.

