## Supplementary figures and images for "Advancing apple genetics research: *Malus coronaria* and *Malus ioensis* genomes and a gene family-based pangenome of native North American apples"

### Fig S1 contact maps.png

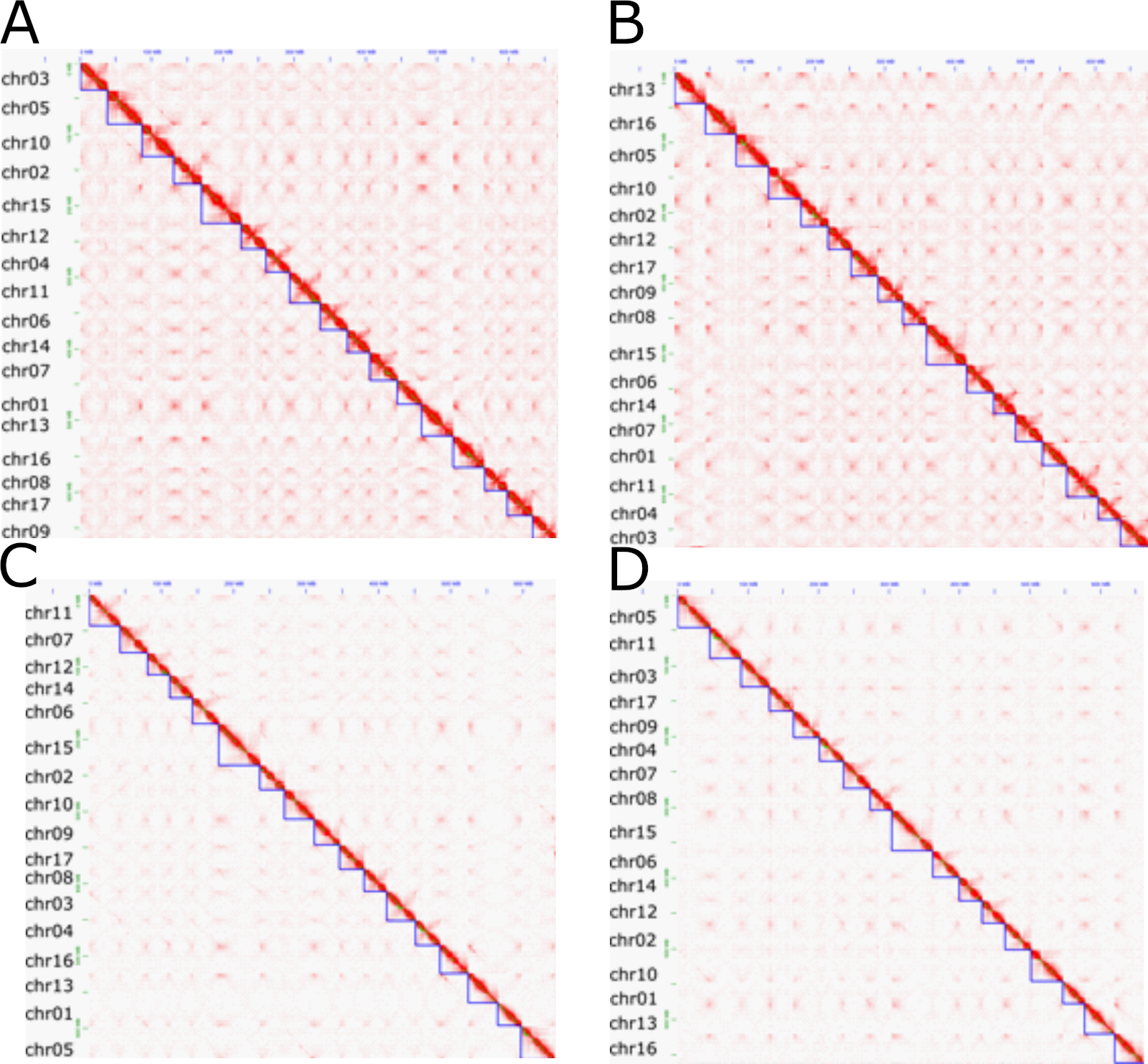

### Fig S2 dotplots.png

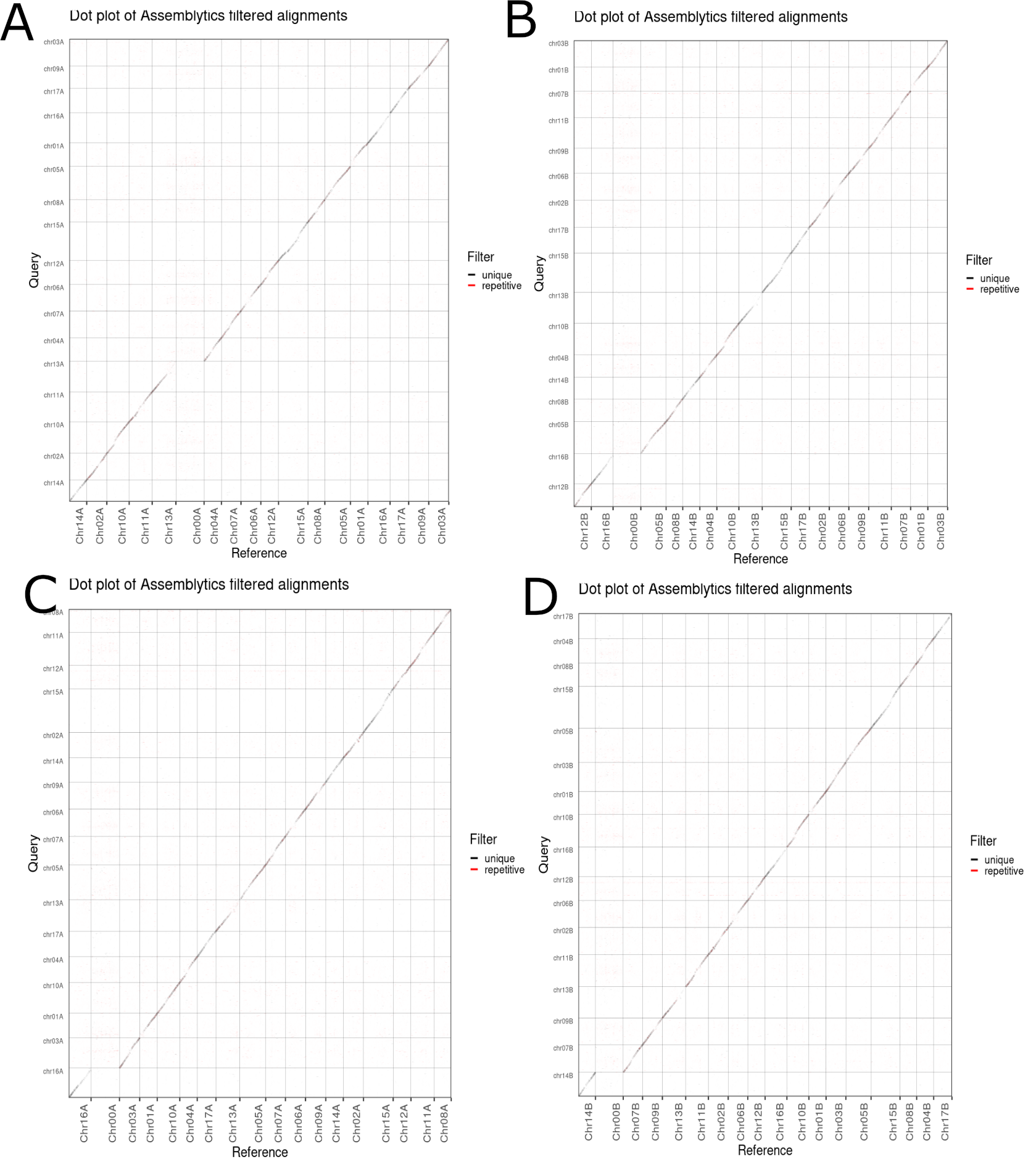

### Fig S3 GE_enrichment_species_fdr_cutoff_corrected.png

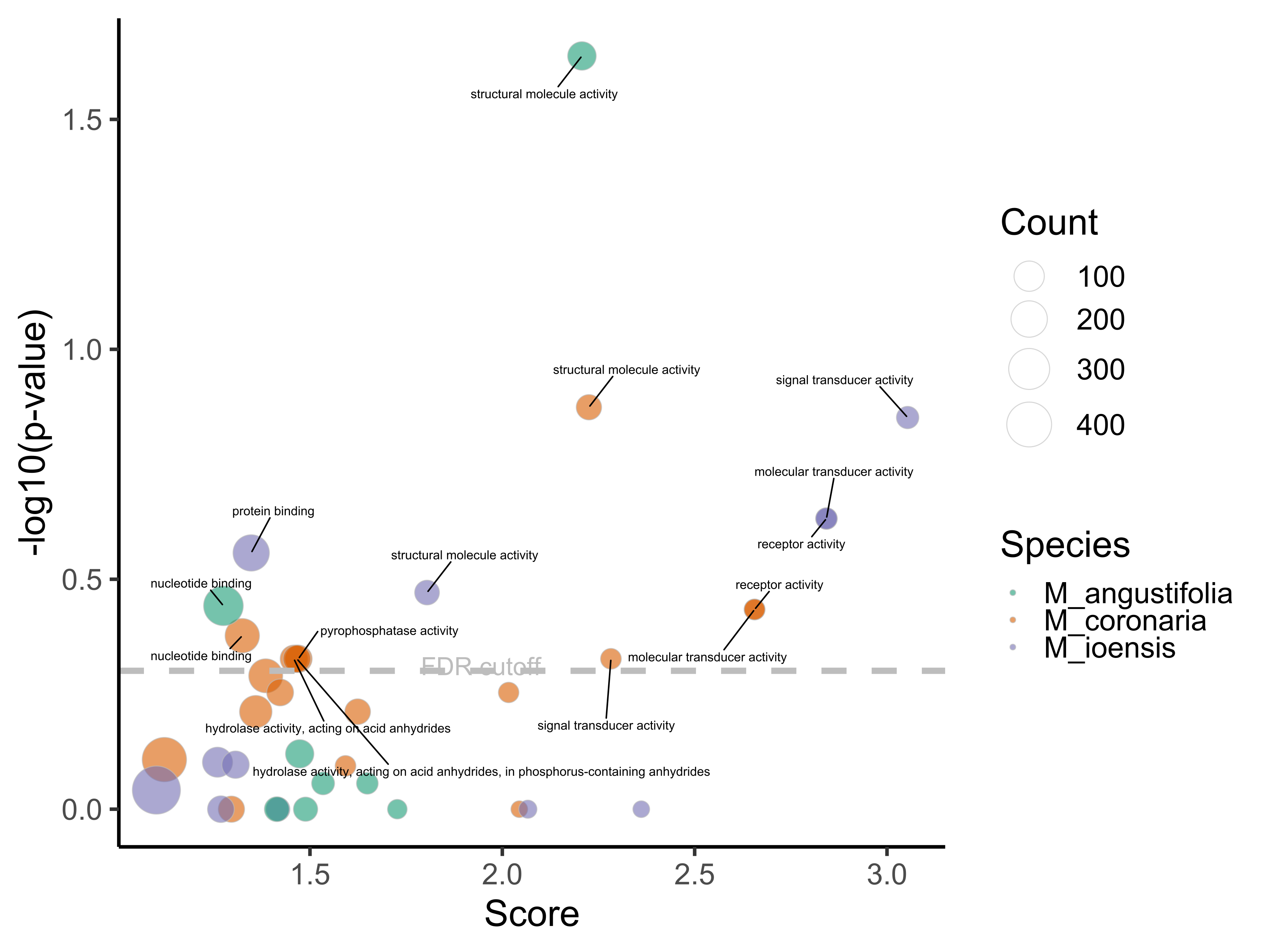

### Fig S4 svs.png

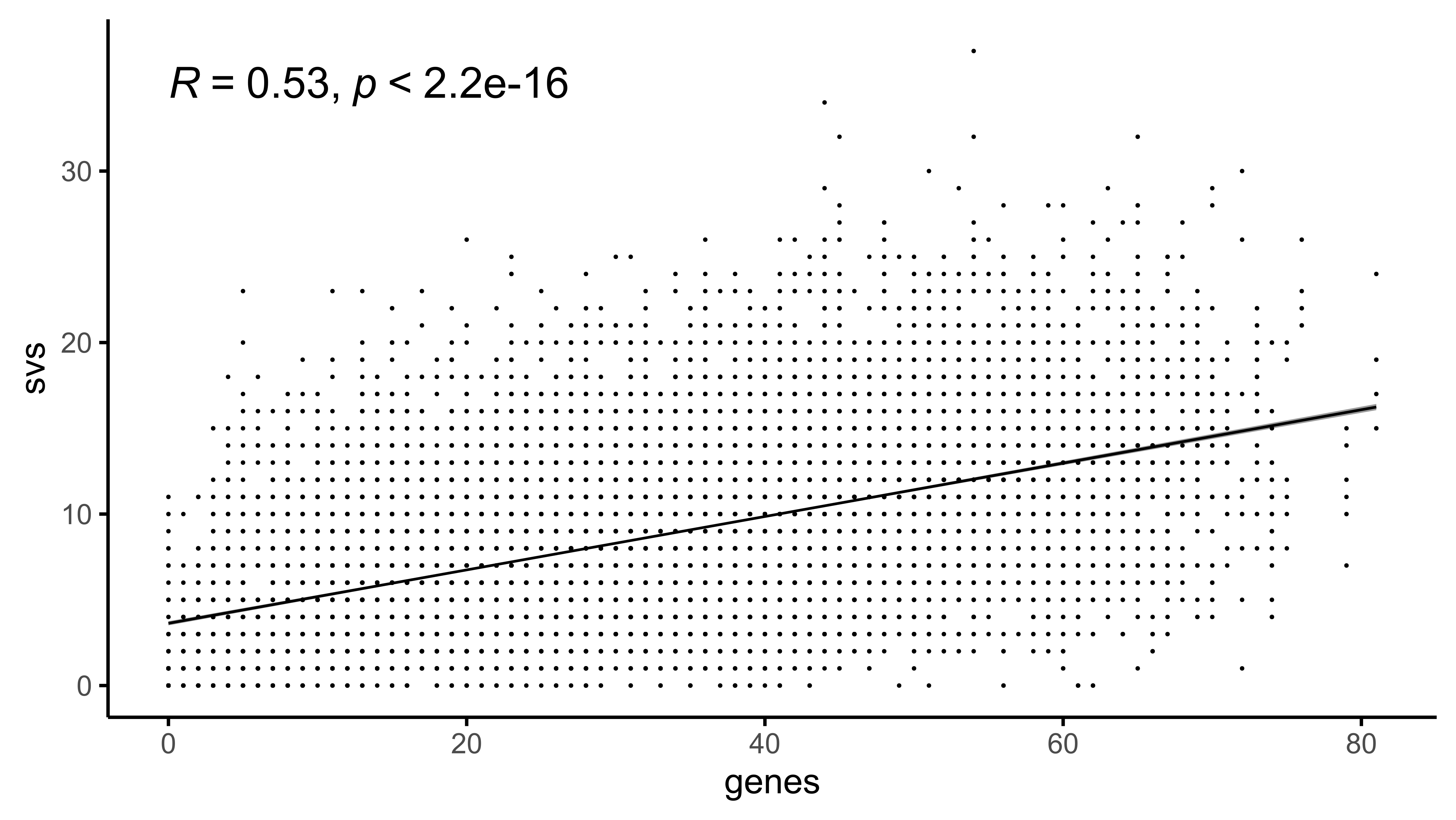
